# Deep dive into the diversity and properties of rhodopsins in actinomycetes of the family *Geodermatophilaceae*

**DOI:** 10.1101/2024.05.16.594530

**Authors:** Sergey V. Tarlachkov, Irina P. Starodumova, Olga V. Boueva, Sergei V. Chernyshov, Lyudmila I. Evtushenko

## Abstract

Rhodopsin with the NTQ motif and heliorhodopsins, both discovered for the first time in actinomycetes of the family *Geodermatophilaceae* were characterized, along with characterization of the DTEW, DTEF and NDQ rhodopsins reported in members of this family previously. The identified absorption maxima ranged from 513 to 559 nm, of which the blue-shifted rhodopsin contained the NTQ motif and the red-shifted one were heliorhodopsin. An assessment of the pumping activity of the NTQ rhodopsin and heliorhodopsin showed that the former functioned as inward Cl^-^-, Br^-^-, I^-^-pump with much weaker activity towards NO^3-^, while no any activity was detected for the later. The DTEW and DTEF rhodopsins had the outward H^+^-transport activity which was dependent on the presence of Ca^2+^ ions and on the particular *E. coli* expression strain. The outward pumping of H^+^ was also detected for NDQ rhodopsin both in NaCl and KCl solutions at pH 5 and 6, but not at pH 7. Weak Na^+^-efflux activity for this rhodopsin was observed at pH 6 and 7 in NaCl solution only after adding carbonyl cyanide m-chlorophenyl hydrazone (CCCP).

## INTRODUCTION

Rhodopsins are a large family of integral membrane proteins containing retinal chromophores that perform the photoreceptor function and also the function of ion pumps in the processes of light energy conversion. Rhodopsins are classified into Type I microbial and Type II animal rhodopsins (play a key role in animal vision). Microbial and animal rhodopsins contain all-*trans* and 11-*cis* retinal molecules, respectively, that capture light (Rozenberg et al., 2021). Recently, a new family of rhodopsins, heliorhodopsins (HeR), was discovered by the functional metagenome analysis, and their global distribution in Archaea, Bacteria, Eukarya and viruses was shown (Pushkarev et al., 2018; Shihoya et al., 2019; Hososhima et al., 2022). Heliorhodopsins are distant relatives of Type I microbial rhodopsins, and their most distinct difference from conventional rhodopsins is inverted membrane topology (Pushkarev et al., 2018; Kovalev et al., 2020; Shihoya et al., 2019).

It is worth noting that most studies within the last two decades have focused on discovery and characterization of various microbial rhodopsins from marine and freshwater microorganisms (Chazan et al., 2022; Fuhrman et al., 2008; Chuon et al., 2021; Keffer et al., 2015; Nakamura et al., 2016; Sharma et al., 2008; Dwulit-Smith et al., 2018; Pushkarev and Béjà, 2016; Pushkarev et al., 2018; Hososhima et al., 2022; Singh et al., 2023). Little is known about distribution and the biological role of these proteins in microorganisms inhabiting terrestrial environments, especially extreme ones. By metagenomic approaches the rhodopsins were revealed, for instance, in phyllospheres of plants, soil crust and Antarctic Dry Valley edaphic systems (Atamna-Ismaeel et al., 2012; Finkel et al., 2013; Guerrero et al., 2017). Rhodopsins were also found in some cultured prokaryotes from terrestrial ecosystems, such as *Exiguobacterium sibiricum* from Siberian permafrost (Petrovskaya et al., 2010; Dioumaev et al., 2013), the thylakoid-lacking cyanobacterium *Gloeobacter violaceus* from calcareous rock (Choi et al., 2014), and *Sphingomonas paucimobilis* from hospital respirator (Maliar et al., 2020). The rhodopsins from the aforementioned organisms have been well characterized and shown to possess H^+^-pumping activity. A Cl^-^-pumping rhodopsin was found in cyanobacterium *Mastigocladopsis repens*, a soil isolate (Hasemi et al., 2016).

Recently, the presence of rhodopsins was reported in actinomycetes (formerly called actinobacteria) of the family *Geodermatophilaceae* (class *Actinomycetes*, phylum *Actinomycetota*) which originate primarily from hot and arid terrestrial environments (Tarlachkov et al., 2020; Tarlachkov et al., 2023). By using a bioinformatics-based approach, the rhodopsins with DTE and NDQ amino acid motifs in positions 85, 89 and 96 (in BR numbering) were discovered in members of this family. The cluster analysis was further showed that the DTE rhodopsin group consisted of two separate groups, DTEW and DTEF, which differed in the presence of highly conserved tryptophan (DTEW) and phenylalanine (DTEF) at position 182 (in BR numbering). Thus, three different groups of microbial rhodopsins were identified in *Geodermatophilaceae* before the present work, namely, DTEW, DTEF and NDQ (Tarlachkov et al., 2020; Tarlachkov et al., 2023).

The DTEW rhodopsins are well known and occur in various groups of microorganisms, including actinomycetes (Chuon et al., 2021; Keffer et al., 2015; Nakamura et al., 2016). The DTEF rhodopsins, on the contrary, are quite unique and have been found so far only in representatives of *Geodermatophilaceae*. As for the rhodopsins with NDQ motif, these have not been described in the literature among actinomycetes and other gram-positives (monoderms), except for a representative of *Geodermatophilaceae* (Tarlachkov et al., 2020), although some close protein sequences of actinomycete origin are present in public protein databases. However, the physiological role and functional characteristics of rhodopsins from *Geodermatophilaceae* have not been studied.

Although generally considered endemic to soils, evolution of members of *Geodermatophilaceae* has continued in specialized biotopes, including rocks and desert sandy soils in areas of moisture deficiency, low nutrients, high salinity and high UV- and gamma-radiation (Busarakam et al., 2016; Tarlachkov et al., 2023; Urzì et al., 2001). The rhodopsin-containing *Geodermatophilaceae* inhabiting such specialized land biotopes represent unique group for study and broadening our understanding of the diversity and functions of microbial rhodopsins and their role in biological adaptation to extremal environments.

In this work, we provide new data on the diversity and properties of rhodopsins from actinomycetes of the family *Geodermatophilaceae*.

## MATERIALS AND METHODS

### Bioinformatic analysis

Complete and high-quality draft genome sequences of 110 bacteria of the family *Geodermatophilaceae* were obtained from GenBank, (Supplementary Table S1) and re-annotated using Prokka 1.14.6 (Seemann, 2014) with the Barrnap 0.9 plug-in. In order to search and identify the rhodopsins genes present in the genomes studied, the reference protein sequences of well-known rhodopsins were used as a query in BLASTp search against translated CDS which were extracted from each genome. A best-hits with e-values lower than 0.01 were considered as potential rhodopsin proteins. Then, each found sequence was aligned using MUSCLE (Edgar, 2021) against reference sequences to finally identify rhodopsins.

Multiple sequence alignment of the revealed rhodopsin proteins was performed by MUSCLE. Phylogenetic trees were inferred by MEGA11 (Tamura et al., 2021). The pairwise identity between protein sequences was determined using TaxonDC 1.3 (Tarlachkov and Starodumova, 2017). The sequences closest to the found rhodopsins were identified by BLASTp searches against the NCBI non-redundant protein database (NCBI NR).

### Genome sequencing and assembly

The cells of strain *Geodermatophilus* sp. DSM 44513 were grown at 28 °C for 24 hours on a rotary shaker in a peptone yeast medium containing the following (g/L): peptone, 5; yeast extract, 3; K_2_HPO_4_, 0.2; glucose, 5; NaCl, 3; pH 7. To obtain long-reads, DNA was isolated with GeneJET Genomic DNA Purification Kit (Thermo Scientific, USA). To enhance cells lysis, enzymatic cell wall digestion was performed in buffer containing 25 mM Tris-HCl 8.0, 0.05 mM EDTA, 40 mM NaCl, 0.1% Triton X-100, and supplemented with 1 mg/ml lysozyme and 0.09 mg/ml EndoRB49 (Chernyshov et. al, 2023). Library were prepared using Ligation Sequencing Kit (Oxford Nanopore Technologies, UK) and Native Barcoding Expansion 13-24 (Oxford Nanopore Technologies, UK). The resulting library was sequenced with MinION flow cell R9.4.1 (Oxford Nanopore Technologies, UK).

For obtaining short-reads, DNA isolation was performed using a FastDNA SPIN Kit and FastPrep-24 homogenizer (MP Biomedicals, USA). Library was prepared with a KAPA HyperPlus Kit (KAPA, USA) and sequenced on an Illumina NovaSeq 6000 platform (Illumina, USA) with a NovaSeq 6000 S2 Reagent Kit (200 cycles).

Long reads basecalling was performed using Guppy v6.5.7 (Oxford Nanopore Technologies, UK). Short and low-quality reads were removed using Filtlong v0.2.1 (https://github.com/rrwick/Filtlong). Genome backbone was assembled using Flye v2.9.2 (Kolmogorov et. al, 2019). Adapter sequences and low-quality regions in short reads were removed using Trimmomatic v0.39 (Bolger et. al, 2014). The assembly was polished with clean short reads using Polypolish v0.5.0 (Wick and Holt, 2022) and Pilon v1.24 (Walker et. al, 2014). The annotation was performed at the NCBI PGAP v6.6 (Tatusova et. al, 2016).

### Rhodopsins cloning and expression

A total of 6 rhodopsin genes were amplified from the genomic DNA of 5 strains (Table 1). The fragments were cloned into the pET-30b expression vector between sites NdeI and XhoI. The termination codon was omitted from all genes, thus cloned genes have LEHHHHHH-coding extension at 3’ end. The gene of DTEF-rhodopsin from *Geodermatophilus obscurus* VKM Ac-658^T^ (GobFR) contains inner XhoI restriction site, which was removed by SOE PCR, along with local optimization of codon usage.

**Table 1.** Rhodopsins characterized in this work and related information. For accession numbers of source sequences, see Supplementary Table S2.

| Rhodopsin designation | Rhodopsin group | Cloned Gene GenBank Acc. number | Strain | Source of strain isolation* |
| --- | --- | --- | --- | --- |
| GdiWR | DTEW | PP765527.1 | *Geodermatophilus dictyosporus* VKM Ac-659<sup>T</sup> (=DSM 43161<sup>T</sup>) | Soil, Westgard Pass, California |
| GdiFR | DTEF | PP765528.1 |  |  |
| GobFR | DTEF | PP765529.1 | *G. obscurus* VKM Ac-658<sup>T</sup> (=DSM 43160<sup>T</sup>) | Soil, Amargosa desert, Nevada |
| MspDR | NDQ | PP765530.1 | *Modestobacter* sp. VKM Ac-2978 | Salt crust on the soil surface, Kysyskum desert, Uzbekistan |
| GspTR | NTQ | PP765531.1 | *Geodermatophilus* sp. DSM 44513 | Marble quarry, Turkey |
| BagHR | HeR | PP765532.1 | *Blastococcus aggregatus* VKM Ac-1606<sup>T</sup> (=DSM 4725<sup>T</sup>) | Sea water, Baltic Sea |

\* Data from Luedemann, 1968; Urzì et al., 2001; Urzì et al. 2004; Tarlachkov et al., 2023.

Protein expression was carried out in *E. coli* BL21(DE3) and C43(DE3) strains using LB medium containing the following (g/L): tryptone, 10; yeast extract, 5; NaCl, 10. Induction was performed with IPTG (isopropyl β-D-1-thiogalactopyranoside) at concentrations of 0.01 mM for *E. coli* BL21(DE3) and 0.5 mM for *E. coli* C43(DE3) for 4 hours at 37 °C with the addition of retinal to a final concentration of 13 μM.

### Protein purification and absorption measurements

All procedures were carried out at 4 °C. Harvested cells were resuspended in a buffer containing 50 mM sodium phosphate, pH 7.8, and 300 mM NaCl, followed by ultrasonication. Undisrupted cells were removed by centrifugation at 5,000g for 10 min. Further, membrane fraction was sedimented at 100,000g for 1 hour, and membrane proteins were extracted using the same buffer supplemented with 1% DDM (n-dodecyl β-D-maltoside). Purification of rhodopsins was carried out on Ni-IDA agarose (Biontex) according to the manufacturer’s recommendations. After elution, buffer was changed to 50 mM Tris-HCl, pH 7.5, 100 mM NaCl, 0.05% DDM using Amicon Ultra Centrifugal Filters. Purity was confirmed by electrophoresis under denaturing conditions. Absorption spectra were recorded with a Shimadzu UV 1800 spectrophotometer.

### Pumping activity assay

Pelleted cells were washed three times with appropriate unbuffered salt solution at a concentration of 100 mM, and then re-suspended for ion transport measurements. The sample was illuminated by 500W halogen lamp supplemented with IR filter. The pH changes were measured in the thermostated cell at 20 °C with intensive stirring and recorded in time-dependent manner. CCCP (carbonyl cyanide m-chlorophenyl hydrazone) at final concentration of 30 μM was used to verify and discriminate proton and non-proton pumping activities.

## RESULTS AND DISCUSSION

### New groups of rhodopsins

As mentioned previously, three groups of rhodopsins with DTEW, DTEF, and NDQ motifs were found in members of the family *Geodermatophilaceae* by bioinformatic analysis (Tarlachkov et al., 2020). A more extended analysis of the available genome sequences of 110 members of this family, including *Geodermatophilus* sp. DSM 44513, sequenced in this work, revealed 69 strains containing the aforementioned DTEW, DTEF, and NDQ rhodopsins, and also NTQ rhodopsin (presumably Cl^-^ pump) and heliorhodopsins. Thus, five groups of rhodopsins have been discovered in members of *Geodermatophilaceae* to date (Fig. 1, Supplementary Table S1, S2).

**Fig. 1.**
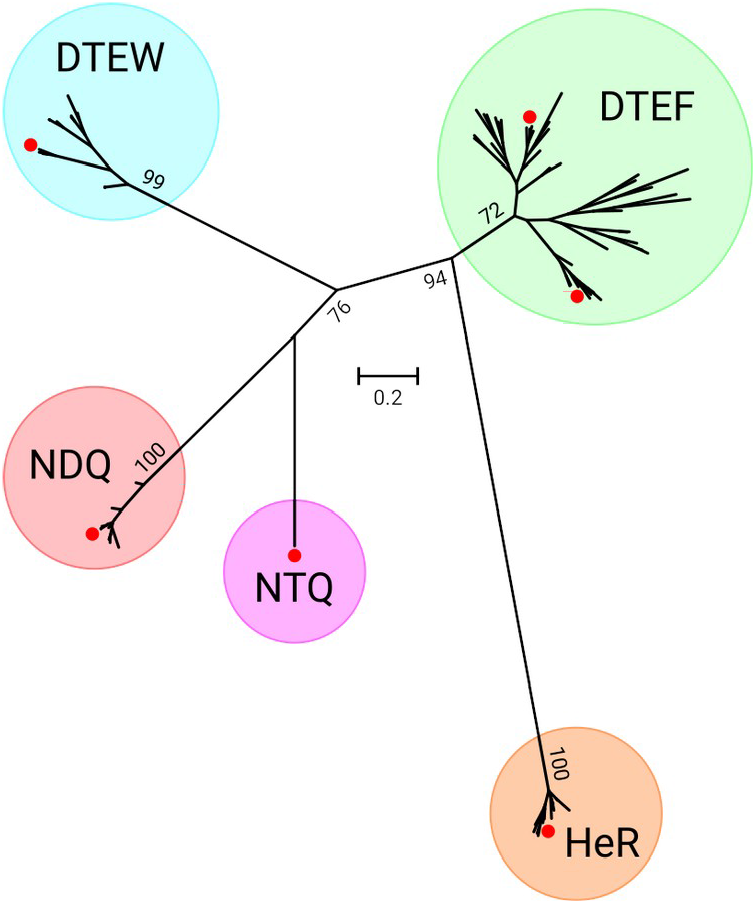
Unrooted approximate maximum likelihood tree of amino acid sequences of rhodopsins from actinomycetes of the family *Geodermatophilaceae*. Red dots denote positions of rhodopsins characterized in this work.

### General characteristics of the genome of *Geodermatophilus* sp. DSM 44513

A complete genome of *Geodermatophilus* sp. DSM 44513, an isolate from marble quarry, Turkey (Urzì et al., 2001) was sequenced and submitted to GenBank under accession number CP135963.1. The genome consisted of a 4.6 Mb circular chromosome with 75,17% G+C content and had a total of 4,454 predicted genes, of which 4,350 were protein-coding. The search revealed genes encoding rhodopsin with DTEW motif and also two rhodopsins novel for the family *Geodermatophilaceae*, NTQ rhodopsin and heliorhodopsin.

### NTQ rhodopsin

The aforementioned novel rhodopsin with a NTQ motif at positions 85, 89 and 96 (in BR numbering) from *Geodermatophilus* sp. DSM 44513 (GspTR) formed a distinct branch on the unrooted tree of rhodopsins found in representatives of the family *Geodermatophilaceae* (Fig. 1). This rhodopsin also clustered apart from the clades of previously reported NTQ rhodopsins acting primarily as Cl^-^ pumps (Supplementary Fig. S3).

To date, only three NTQ rhodopsins have been characterized in detail, namely, NM-R3 from *Nonlabens marinus* (*Flavobacteriia*) (Yoshizawa et al., 2014), FR from *Fulvimarina pelagi* (*Alphaproteobacteria)* (Inoue et al., 2014) and AaClR from *Alteribacter aurantiacus* (*Bacillaceae*) (Singh et al., 2023). In addition, two identified but incompletely characterized rhodopsins from this group have been reported: PoClR from *Parvularcula oceani* (*Alphaproteobacteria*) (Inoue et al., 2016) and unnamed protein from *Nonlabens* sp. YIK11 (*Flavobacteriia*) (Kwon et al., 2016). There is also information on some uncharacterized NTQ rhodopsins originating from marine bacteria (Yoshizawa et al., 2014, Kwon et al., 2016). All the above bacteria with NTQ rhodopsins, except for *Alteribacter aurantiacus* and *Geodermatophilus* sp. DSM 44513, are Gram-negatives (diderms), and all originate from aquatic, mainly marine, environments, with the exception of the above *Geodermatophilus* sp. DSM 44513 isolated from a terrestrial source (marble quarry) (Urzì et al., 2001).

In the NCBI protein database, the sequences nearest to the NTQ rhodopsin from *Geodermatophilus* sp. DSM 44513 (GspTR) were found in uncultivated actinomycetes, such as a representative of the suborder *Frankineae* from biological soil crust sampled in Negev Desert (GenBank acc. CAA9312725.1, 79.6% identity) and a member of the order *Mycobacteriales* from soil (GenBank acc. HWG93181.1, 63.0% identity). Other found rhodopsins showed less than 43% identities to GspTR and did not always contain NTQ motif. Thus, the phylogenetically distant GspTR rhodopsin discovered in *Geodermatophilus* sp. DSM 44513 and some related satellite sequences from uncultivated organisms available in the NCBI protein database form a separate group of hypothetical Cl^-^ pumps which appear to specialize for non-aquatic ecological niches.

GspTR rhodopsin and its homologs from the two above non-aquatic actinomycetes, members of *Frankineae* and *Mycobacteriales*, shared less than 38.9% amino acid identity to the all other described NTQ rhodopsins from aquatic bacteria and had several noticeable differences in key amino acids. For example, GspTR contained a hydrophobic valine at position 204 (in BR numbering), while the NTQ rhodopsins from the above aquatic bacteria had a charged arginine at homologous positions (Supplementary Fig. S4). As a result of such substitution, the H^+^ release mechanisms may alter due to the possible involvement of the corresponding amino acid residues in the proton-releasing group (PRG).

### Heliorhodopsins

A total of 18 heliorhodopsin genes were discovered in actinomycetes of the family *Geodermatophilaceae* by the bioinformatic analysis (Supplementary Table S1, S2). These were present predominantly in strains of the genus *Modestobacter* and also identified in three *Blastococcus* strains and three representatives of *Geodermatophilus*. The detected heliorhodopsins were assigned to subfamily 1 according to the classification of Kovalev et al. (Kovalev et al., 2020) and to group A according to Chazan et al. (Chazan et al., 2022). The subfamily 1 and group A contain mostly proteins from actinomycetes, including HeR 48C12, a well-studied member of the heliorhodopsin family found in an actinomycete from the freshwater Lake Kinneret (Pushkarev et al., 2018).

The identified *Geodermatophilaceae* heliorhodopsins often co-occurred with NDQ rhodopsins in the same strains (Supplementary Table S1, S2). In two strains, *M. roseus* DSM 45764 and *M. marinus* DSM 45201, the genes for heliorhodopsins and NDQ-rhodopsins were closely located, but did not form an operon (Supplementary Fig. S5). The heliorhodopsin genes in *Blastococcus* sp. DSM 46792, *Modestobacter* sp. VKM Ac-2978 and *M. caceresii* KNN 45-2b were frame-shifted or fragmented (Supplementary Fig. S6), in contrast to all other rhodopsin genes in members of the family *Geodermatophilaceae*, and are most likely inactive. The above findings suggest a weaker selection pressure for the heliorhodopsin genes as compared with the other rhodopsins, at least in *Geodermatophilaceae*. On the other hand, the heliorhodopsin proteins of *Geodermatophilaceae* have a relatively large percentage of identity among themselves, typically more than 85%, which may point to some conservation processes.

Heliorhodopsins from the family *Geodermatophilaceae* showed less than 49% identity with HeR 48C12, the aforementioned well-studied heliorhodopsin from actinomycete (Pushkarev et al., 2018). But at the same time, the *Geodermatophilaceae* heliorhodopsins have conserved most of the reported functionally important amino acids (Supplementary Fig. S7). Notable differences in amino acid sequences were revealed at positions associated with dimerization and the AB loop, and also with serine-to-alanine substitution at position 76 (in 48C12 numbering), which is part of the retinal Schiff base (RSB).

To date, there is no consensus on the biological and functional roles of heliorhodopsins. One of the approaches to understanding their role is the analysis of neighboring genes. From the strongest association between heliorhodopsins and the DegV family proteins involved in activation of fatty acids, it was hypothesized that heliorhodopsins might be involved in light-induces membrane lipid modifications (Chazan et al., 2022). However, genes for DegV proteins were not found in the vicinity of heliorhodopsin genes in strains of *Geodermatophilaceae* (Supplementary Fig. S5), although the DegV genes were present in the majority of members of this family, including the heliorhodopsin-lacking ones. From the above findings it follows that if any relationship between heliorhodopsins and DegV takes place, it is not universal.

The heliorhodopsins were also reported to regulate the activity of class 2 cyclobutane pyrimidine dimer photolyase (Shim et al., 2022) and glutamine synthetase (Cho et al., 2022). In both cases, the genes for the target enzymes were adjacent and formed an operon with heliorhodopsins. It has also been noted that the heliorhodopsin genes from some aquatic actinomycetes (actinobacteria in the original work) are surrounded by two large clusters of Nuo genes, the products of which are the key proteins in respiratory chains (Kovalev et al., 2020). A different situation was observed in *Geodermatophilaceae*, where the heliorhodopsin genes had a diverse vicinity without clear patterns. In addition, they were predominantly oriented in the opposite direction to the nearest neighboring genes. The above data provide clear evidence for the absence of co-expression. Thus, it can be stated that the analysis of neighboring genes does not allow us to make an assumption about functions of heliorhodopsins in the *Geodermatophilaceae* family.

### Light-harvesting carotenoid antenna

A previous study of *Salinibacter ruber* (*Actinomycetales*) has shown that it produces xanthorhodopsin (XR), a well-known light-driven H^+^ pump (Balashov et al., 2005). In addition to retinal, XR contains a non-covalently bound carotenoid salinixanthin with a function of a light-harvesting antenna (Balashov et al., 2005). In the present study, we have found that a significant number of amino acids involved in carotenoid binding by xanthorhodopsin are present in Type I rhodopsins from *Geodermatophilaceae*, including glycine at a position homologous to 156 in xanthorhodopsin (Supplementary Fig. S8). Glycine at this position was supposed to be the best diagnostic feature for the possibility of binding the salinixanthin-like carotenoids (Luecke et al., 2008).

The Na^+^-pumping rhodopsin (NaR) from flavobacterium *Dokdonia* sp. PRO95 was reported to bind carotenoid antenna after replacement of threonine by glycine at position 216 (homolog of G201 for xanthorhodopsin) (Anashkin et al., 2018). Interestingly, glycine at the corresponding position was found in most Type I rhodopsins from *Geodermatophilaceae*, and only NDQ rhodopsins had methionine instead of glycine. Other carotenoid-binding amino acids identical between xanthorhodopsin and NaR rhodopsin from *Dokdonia* sp. PRO95 (G156, N191, L197, Y207 in XR numbering) were the same as in the majority of Type I rhodopsins of *Geodermatophilaceae*. From all the above data it might be supposed that the DTE and NTQ rhodopsins from *Geodermatophilaceae* may bind the light harvesting antenna and enhance efficiency of ion transport, like that reported for other carotenoid-binding rhodopsins.

### Rhodopsins absorption spectra

DDM-solubilized rhodopsins isolated from *E. coli* were violet to red in color, with absorption maxima ranging from 513 to 559 nm at pH 7.5 (Fig. 2). The NTQ rhodopsin from *Geodermatophilus* sp. DSM 44513 (GspTR) had absorption peak at 513 nm in 100 mM NaCl and was the most blue-shifted compared to the 24 NTQ rhodopsins available in the Karasuyama et al. database (Karasuyama et al., 2018). The absorption peak of GspTR was the closest to that of FR rhodopsin from *Fulvimarina pelagi* (518 nm in 1660 mM NaCl) (Inoue et al., 2014) and of AaClR from *Alteribacter aurantiacus* (517 nm in 100 mM NaCl) (Singh et al., 2023), and showed significant distance from NM-R3 rhodopsin from *Nonlabens marinus* with absorption peak at 533.5 nm (Yoshizawa et al., 2014).

**Fig. 2.**
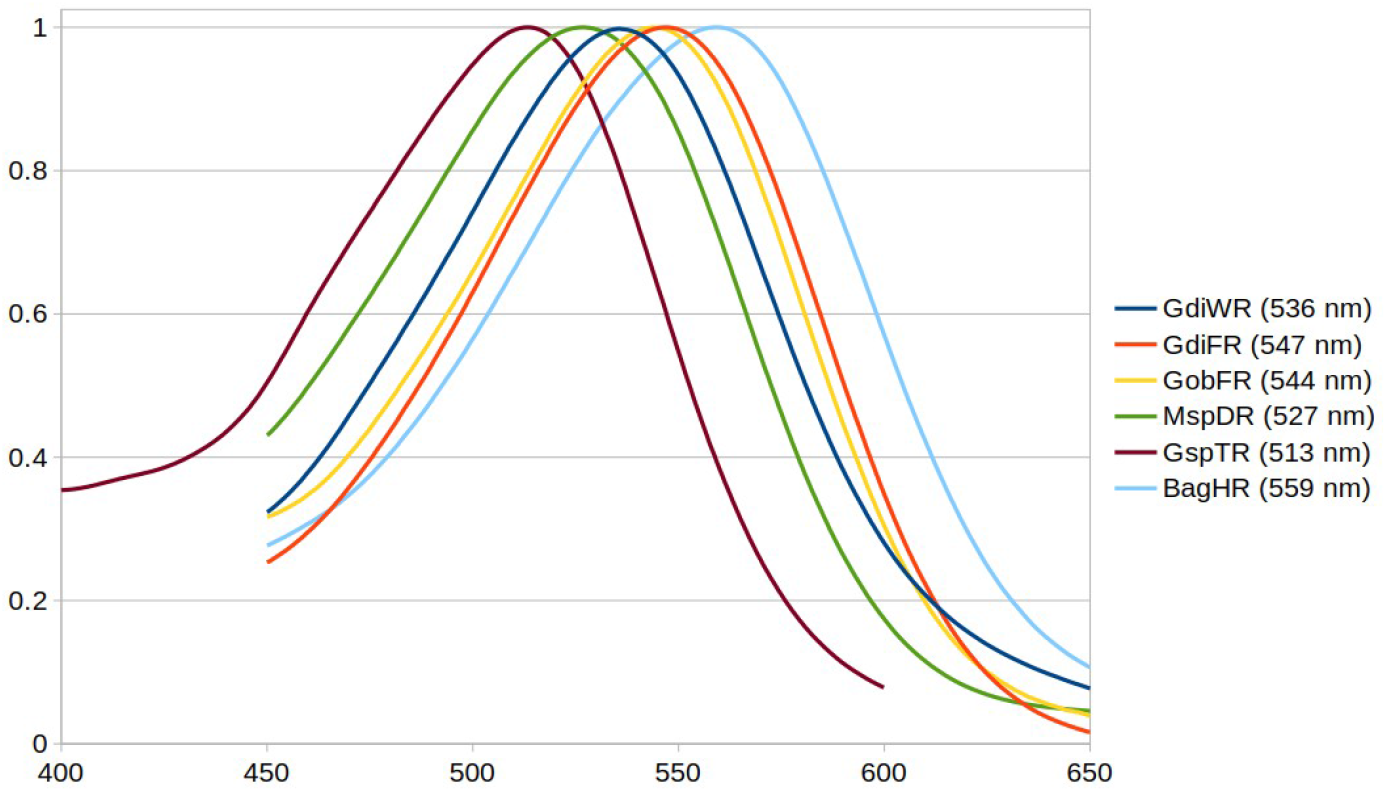
Absorption spectra of purified rhodopsins solubilized in the DDM. For rhodopsins designations, see Table 1.

Heliorhodopsin from *B. aggregatus* VKM Ac-1606^T^ (BagHR) with the absorption maximum at 559 nm was the most red-shifted protein compared to all actinomycete heliorhodopsins and the majority of other heliorhodopsins characterized to date (Kim et al., 2021; Cho et al., 2022; Hososhima et al., 2022; Shihoya et al., 2019; Chazan et al., 2022). This value was only slightly lower than that reported for the known most red-shifted heliorhodopsin HULAa30F3 (absorption peak, 562 nm) from aquatic *Chloroflexi* (Chazan et al., 2022). The *Geodermatophilaceae* heliorhodopsins had less than 43% protein identity to the HULAa30F3 and contained the highly conserved canonical histidine residues at positions 23 and 80 (in 48C12 numbering), unlike hydrophobic leucine in HULAa30F3 (Supplementary Fig. S7). The replacement of histidine with an aromatic phenylalanine at each position was reported to significantly slow down the photocycle of 48C12 (Pushkarev et al., 2018).

### Pumping specificity

#### Rhodopsins with DTEW and DTEF motifs from *G. dictyosporus* and *G. obscurus*

GdiWR and GdiFR rhodopsins from *G. dictyosporus* VKM Ac-659^T^ and GobFR rhodopsin from *G. obscurus* VKM Ac-658^T^ (Table 1) were found to exhibit outward H^+^-pumping activity, which depended on the presence of Ca^2+^ ions and the particular expression strain. In case the rhodopsin genes were expressed in *E. coli* BL21(DE3), the pumping was observed in both CaCl_2_ and NaCl solutions, but the activity was higher at CaCl_2_ (Fig. 3A, 3B, 3G, 3H). On the other hand, Ca^2+^ was strictly required for H^+^-pumping when DTE rhodopsin genes are expressed in strain *E. coli* C43(DE3) (Fig. 3D, 3E, 3J, 3K). Another divalent ion, namely, Mg^2+^, showed little effect only in case the strain *E. coli* BL21(DE3) was used for expression, but no effect was observed with *E. coli* C43(DE3) expression strain (Fig. 3C, 3F, 3I, 3L). The H^+^-pumping activity was confirmed using cells treated with a protonophore (CCCP) that greatly reduced light-induced pH changes.

**Fig. 3.**
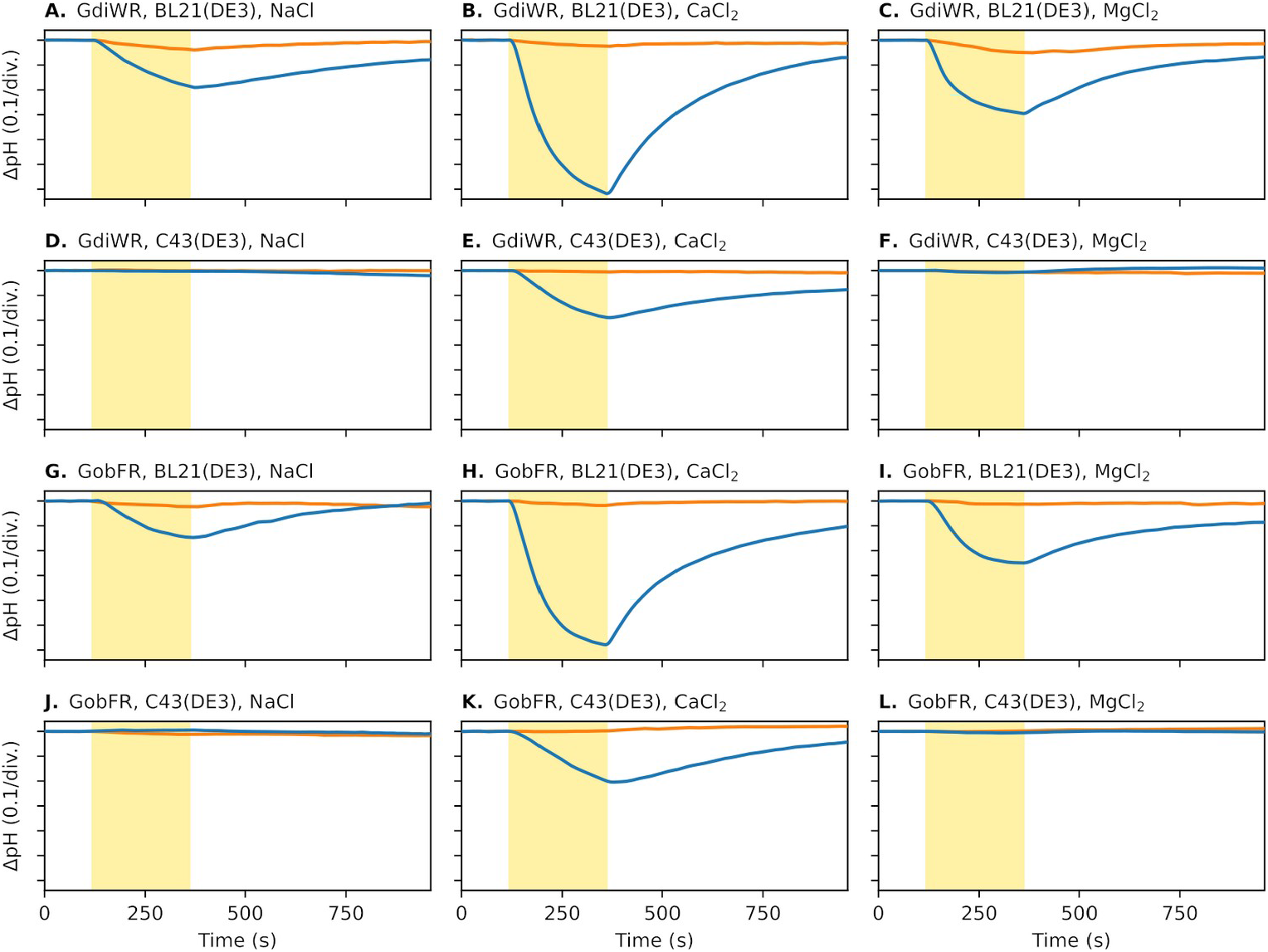
Ion transport activity of the DTE rhodopsins from *G. dictyosporus* VKM Ac-659^T^ (GdiWR) and *G. obscurus* VKM Ac-658^T^ (GobFR) expressed in *E. coli* BL21(DE3) and C43(DE3) strains. For DTEF rhodopsin from *G. dictyosporus* VKM Ac-659^T^ (GdiFR) see Supplementary Fig. S9. Measurements were performed with (orange line) and without (blue line) adding CCCP. The cell suspensions were illuminated for 240 s (yellow area on the plots).

It is worth mentioning that no Ca^2+^ requirement for pumping activities was noted previously for rhodopsins, including those from actinomycetes (Chuon et al., 2021; Dwulit-Smith et al., 2018; Keffer et al., 2015; Nakamura et al., 2016; Pushkarev and Béjà, 2016). However, some papers and guidelines recommend addition of Ca^2+^ and Mg^2+^ to pumping solutions at concentrations of 100 μM and 10 mM, respectively (Han et al., 2020; Maresca et al., 2016; Wang et al., 2003). In our experiments, no DTE rhodopsin activity was observed at the above Ca^2+^ and Mg^2+^concentrations when the expression strain *E. coli* C43(DE3) is used (data not shown). The activity of rhodopsins from other groups studied in this work did not depend on the presence of Ca^2+^ or on the use a particular *E. coli* expression strain.

Considering the requirement for high Ca^2+^ concentrations, it is unlikely that it plays a regulatory function. It can be inferred from our study that Ca^2+^ in a high concentration is necessary to correct folding of DTE rhodopsins. Although the DTE rhodopsins are colored that indicates the overall right protein folding, the local miss-folds may still prevent rhodopsin pumping activity in the absence of Ca^2+^. In addition, Ca^2+^ can interact with lipid membranes, in particular, it rigidifies and orders lipid bilayers by involving in mechanisms of conformational changes of the lipid headgroup region, ordering of acyl chains and lipid dehydration (Melcrová et al., 2016). Thus, Ca^2+^ ions can affect the membrane structure, resulting in the activation of rhodopsins.

It is unclear whether Ca^2+^ is required for pumping process only when the DTE rhodopsins are expressed in Gram-negative (diderm) *E. coli*, or it also necessary in case the expression strain is Gram-positive (monoderm) prokaryotes like the host *Geodermatophilaceae* strains. The Gram-positives do not contain outer membrane, possesses different lipid composition and usually has thicker and more rigid cell walls with multilayered peptidoglycans and different layers of heteropolysaccharides and peptides (Rohde, 2019). On the other hand, the requirement for Ca^2+^ can be unique feature of the DTE rhodopsins from *Geodermatophilaceae* when Ca^2+^ acts as an activator of these rhodopsins under certain physico-chemical environmental conditions, for instance, in the calcium-rich biotopes. This is consistent with the ability of representatives of the family *Geodermatophilaceae* to live in such biotopes, for example, in limestone carbonates. It can also be assumed that, despite the similarity in behavior of DTEW and DTEF rhodopsins when expressed in *E. coli*, their functions may somewhat differ in host strains under certain natural conditions. This supposition follows from the co-presence of rhodopsins from both above groups, DTEW and DTEF, in three members of the family *Geodermatophilaceae* (Supplementary Table S1, S2).

#### Rhodopsin MspDR with NDQ motif from *Modestobacter* sp. VKM Ac-2978

MspDR rhodopsin from *Modestobacter* sp. VKM Ac-2978 acts exclusively as H^+^ pump in both NaCl and KCl solutions at pH 5 (Fig. 4A, 4B). At pH 6, the H^+^-pumping still predominated (Fig. 4C, 4D), but some Na^+^-pumping activity became additionally detectable when CCCP is added to the cell suspension in NaCl solution. Even at pH 7, Na^+^-pumping was observed in NaCl solution only after adding CCCP (Fig. 4E). However, H^+^-pumping was absent at neutral pH according to the experiment with KCl solution (Fig. 4F). As follows, the lack of any activity in NaCl solution at pH 7 without CCCP is not due to a combination of the opposite processes of Na^+^- and H^+^-pumping, just weak activity against Na^+^.

**Fig. 4.**
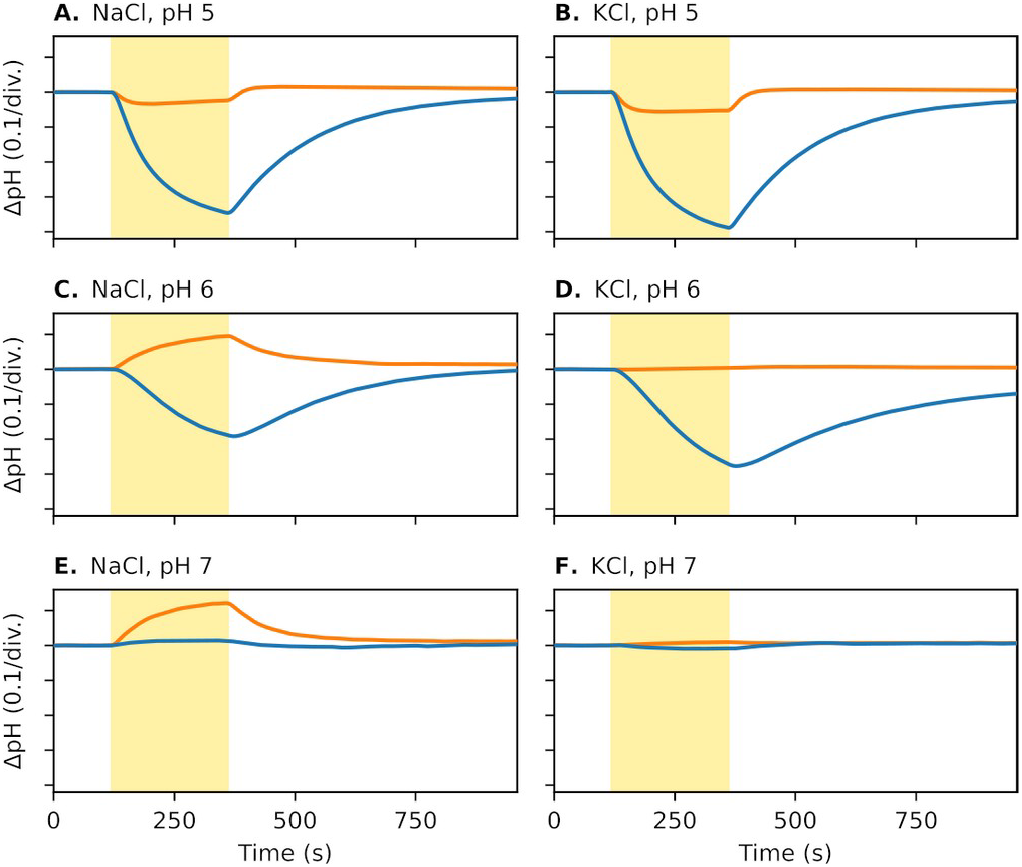
Ion transport activity of NDQ rhodopsin from *Modestobacter* sp. VKM Ac-2978 (MspDR) expressed in *E. coli* C43(DE3). Measurements were performed with (orange line) and without (blue line) adding CCCP. The cell suspensions were illuminated for 240 s (yellow area on the plots).

Previously it was supposed that NDQ rhodopsins from the *Geodermatophilaceae* have Na^+^- or H^+^/Na^+^-pumping activity (Tarlachkov et al., 2020). The present study showed that the MspDR rhodopsin acts predominantly as a pH-dependent H^+^ pump, but the Na^+^ activity in *E. coli* cell suspension was very weak. For strong Na^+^-pumping activity, a much more alkaline pH and much more than 100 mM NaCl concentration are probably required.

Thus, the behavior of MspDR rhodopsin differed from that of the previously characterized NDQ rhodopsins KR2, NdR2 and IAR from *Flavobacteriaceae* and *Cyclobacteriaceae* which showed a significant Na^+^-pumping activity at neutral pH in 100 mM NaCl solution, and H^+^-pumping at any pH in a KCl solution (Inoue et al., 2013; Li et al., 2015; Zhao et al., 2017). MspDR rhodopsin also differed from NM-R2 (Yoshizawa et al., 2014) and GLR (Balashov et al., 2014) rhodopsins found in *Flavobacteriaceae*, which exhibited only Na^+^ activity. It is worth noting that all the aforementioned NDQ rhodopsins are harbored in Gram-negative (diderm) bacteria inhabiting aquatic ecosystems, mainly seas, and their differences in the behavior from MspDR rhodopsin of Gram-positive (monoderm) *Modestobacter* from Kyzylkum Desert can be related to their specific habitat conditions and the cell envelope structure.

Protein sequence analysis revealed also a number of differences in the amino acids between NDQ rhodopsins from *Geodermatophilaceae* and well-studied NDQ rhodopsins from aquatic Gram-negatives (Supplementary Fig. S10). The most notable differences were observed at positions L74, N61 and G263 (in KR2 numbering). The first amino acid residue (L74) plays an important role in switching between Na^+^-to H^+^-pumping activities and the last two (N61 and G263) are associated with ion selectivity (Gushchin et al., 2015; Kovalev et al., 2019). In addition, the valine is detected at the position homologous to H180 in KR2. Histidine at this position was reported to be involved in the mechanism of Na^+^-pumping and is characteristic of all NDQ rhodopsins studied in this respect (Kriebel et al., 2023).

It is clear, that the MspDR rhodopsin and most likely other NDQ rhodopsins from *Geodermatophilaceae* act only as H^+^ pumps at pH < 6. MspDR also showed slight bifunctional properties at pH 6-7, and may probably act as Na^+^ pump in some extreme conditions at high pH and salinity. These rhodopsins are apparently incapable of transporting Na^+^ to a noticeable degree under seawater conditions, but they can do this in case the Na^+^ content and pH value of the environment are significantly higher than in seawater (>>0.5 M NaCl and pH >>8). This may be important for the survival and vital activity of *Geodermatophilaceae* species in saline and hyper-arid environments under the salinity level much higher than in seawater (Busarakam et al., 2016; Tarlachkov et al., 2023).

On the other hand, members of the family *Geodermatophilaceae* with NDQ rhodopsins can grow well in the NaCl-lacking nutrient media (for example, ISP2) (Busarakam et al., 2016; Tarlachkov et al., 2023; Xiao et al., 2011), unlike most marine microorganisms with Na^+^- or Na^+^/H^+^-pumping rhodopsins that prefer salt-containing media (Khan et al., 2006; Park et al., 2012; Yoon et al., 2006). It can be therefore supposed that the NDQ rhodopsins revealed in *Geodermatophilaceae* are capable of performing two functions, namely, to maintain the Na^+^ homeostasis under extreme salinity and the pH balance under acidification of the environment.

Previously a pH-sensitive DTS rhodopsin active at pH 6 and below from Gram-negative (diderm) bacterium *Sphingomonas paucimobilis* promising for use in optogenetics has been reported (Maliar et al., 2020). Microbial rhodopsins with the NDQ motif from actinomycetes of the family *Geodermatophilaceae* may represent another group of pH-sensitive rhodopsins of interest for optogenetic studies and applications. Further improvement in specificity can be achieved, in particular, by site-directed mutagenesis to completely eliminate the affinity to Na^+^ ions. These proteins can also be used as model to study the mechanism of switching between the H^+^ and Na^+^ specificities in NDQ rhodopsins.

#### Rhodopsin GspTR with NTQ motif from *Geodermatophilus* sp. DSM 44513

GspTR rhodopsin from *Geodermatophilus* sp. DSM 44513 showed high inward Cl^-^- and Br^-^-pumping activities (Fig. 5A, 5B, 5C, 5D), and had weaker activity towards I^-^ (Fig. 5E, 5F). Some trace signals were also registered in pumping solutions with NO_3_ ^-^ (Fig. 5G, 5H), but no any pumping activity against SO_4_ ^2-^ was detected (Fig. 5I, 5J). It was observed during experiment that even at a slight contamination of the cell suspension with Cl^-^ ions a noticeable signal was detected, so a double junction electrode filled with KNO_3_ was used to prevent such contamination from the reference electrode.

**Fig. 5.**
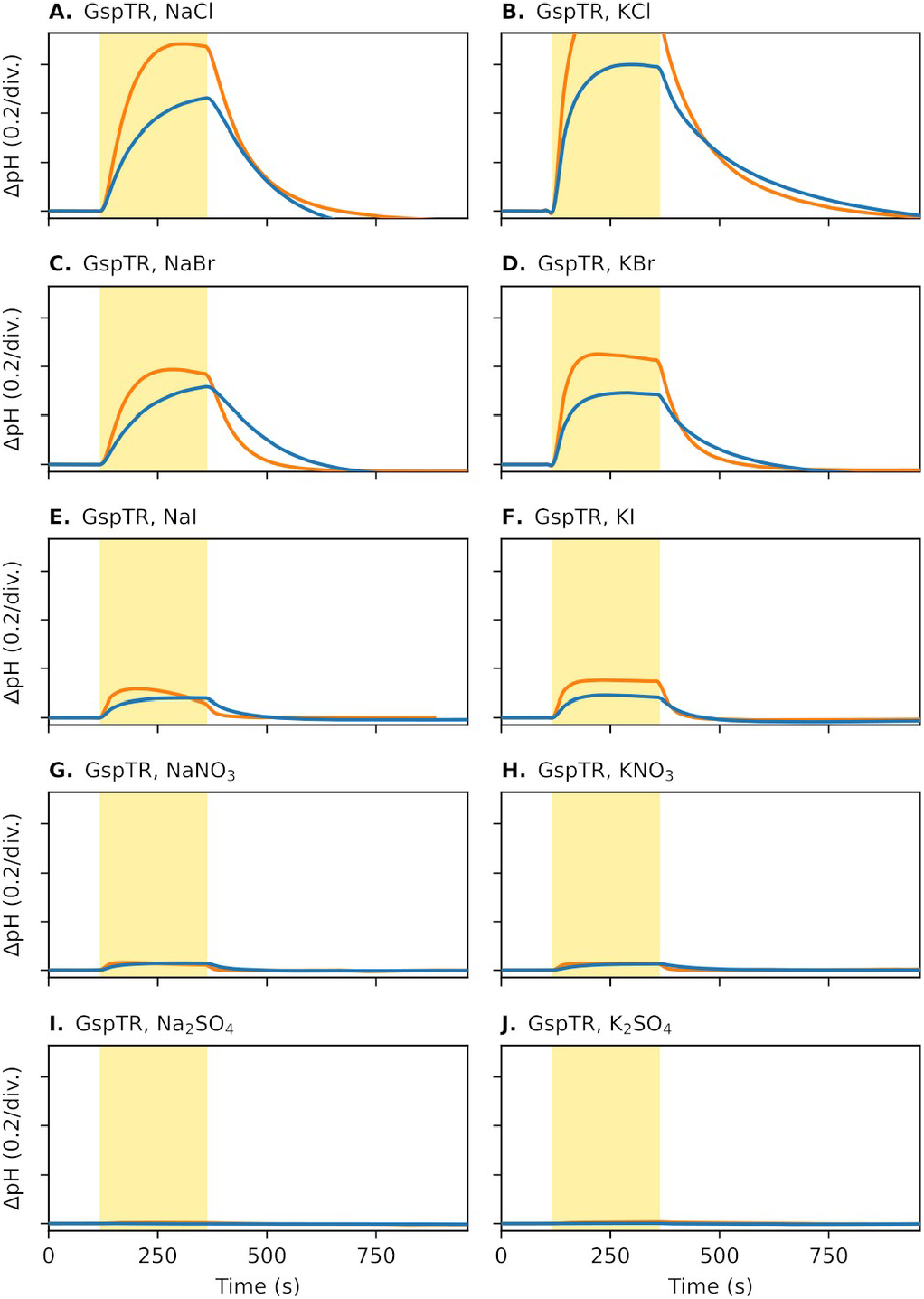
Ion transport activity of NTQ rhodopsin from *Geodermatophilus* sp. DSM 44513 (GspTR) expressed in *E. coli* C43(DE3). Measurements were performed with (orange line) and without (blue line) adding CCCP. The cell suspensions were illuminated for 240 s (yellow area on the plots).

#### Heliorhodopsin BagHR from *B. aggregatus*

BagHR from *B. aggregatus* VKM Ac-1606^T^ did not exhibit any pumping activity in either NaCl or CaCl_2_ solutions (data not shown). Some factors may interfere with the pumping activity of this heliorhodopsin, like that described above for DTE rhodopsins. Thus, the functions of heliorhodopsin from *B. aggregatus* (and also from other members of *Geodermatophilaceae*) remain unclear. On the other hand, these functions may have no relation to pumping process, and the *Geodermatophilaceae* heliorhodopsins may play completely different functional and biological roles. Diverse biological functions of heliorhodopsins were supposed based on their 10 different groups (subfamilies) identified by bioinformatic analysis (Kovalev et al., 2020) and also follows from the data discussed above in the heliorhodopsin-related section. As a result, it can be assumed that taxonomic or ecological groups of the family *Geodermatophilaceae* may have their own specific heliorhodopsins with significantly different functions acquired as a result of their evolution and adaptation to specific terrestrial ecological niches.

## CONCLUSION

New data have been provided on the diversity of rhodopsins in actinomycetes of the family *Geodermatophilaceae* (class *Actinomycetes*, phylum *Actinomycetota*) mostly originating from hot and arid ecosystems, and some new properties and functional characteristics of the rhodopsins have been reported. Rhodopsins from all five groups studied in the present work, both the newly discovered NTQ and heliorhodopsins, and the DTEW, DTEF and NDQ rhodopsins previously identified in the *Geodermatophilaceae*, have either unique absorbance maxima or unique features of pumping activity. The retinal proteins of Gram-positive (diderm) *Geodermatophilaceae* share quite low similarities to the respective rhodopsins from Gram-negatives (monoderms) inhabiting marine and freshwater environments and also differ from them in some highly conserved amino acid residues. This is not surprising given the evolutionary distances and phenotypic differences between rhodopsin-producing members of *Actinomycetota* and representatives of the other bacterial phyla, as well as remarkable variations between their specialized biotopes, both aquatic and terrestrial, including rocks and desert sandy soils. Further study will extend our knowledge about the diversity and functional characteristics of the *Geodermatophilaceae* rhodopsins and their role in adaptation of actinomycetes of this family to harsh environments.

## ABBREVIATIONS

AaClR: *Alteribacter aurantiacus* rhodopsin
BR: bacteriorhodopsin
CCCP: carbonyl cyanide m-chlorophenyl hydrazone
CDS: coding sequence
DDM: n-dodecyl β-D-maltoside
FR: *Fulvimarina pelagi* rhodopsin
GLR: *Gillisia limnaea* rhodopsin
HeR: heliorhodopsins
IAR: *Indibacter alkaliphilus* rhodopsin
IPTG: isopropyl β-D-1-thiogalactopyranoside
ISP2: International *Streptomyces* Project-2
KR2: *Krokinobacter eikastus* (*Dokdonia eikasta*) rhodopsin 2
LB: lysogeny broth
NdR2: *Nonlabens dok-donensis* rhodopsin 2
NM-R2: *Nonlabens marinus* rhodopsin 2
NM-R3: *Nonlabens marinus* rhodopsin 3
PCR: polymerase chain reaction
PoClR: *Parvularcula oceani* rhodopsin
SOE PCR: splicing by overhang extension PCR
XR: xanthorhodopsin.

## ACKNOWLEDGEMENTS

This work was financially supported by the Ministry of Science and Higher Education of the Russian Federation (Grant agreement # 075-15-2021-1051).

The authors are grateful to prof. Oded Béjà for pointing out the presence of heliorhodopsins in the *Geodermatophilus* family.

