## Supplementary for "Deep dive into the diversity and properties of rhodopsins in actinomycetes of the family *Geodermatophilaceae*"

**Supplementary Table S1.** The *Geodermatophilaceae* genomes analyzed in the present study and the revealed rhodopsins.

| # | Assembly Acc. No. | Organism | Rhodopsin(s)[1] |
|---|---|---|---|
| 1 | GCA_900221005.1 | *Blastococcus aggregatus* DSM 4725$^T$ | HeR |
| 2 | GCA_002938425.1 | *Blastococcus atacamensis* P6$^T$ | DTEF |
| 3 | GCA_900102425.1 | *Blastococcus aurantiacus* DSM 44268$^T$ | DTEF |
| 4 | GCA_030165005.1 | *Blastococcus capsensis* BMG 804$^T$ | NDQ |
| 5 | GCA_006717095.1 | *Blastococcus colisei* DSM 46837$^T$ | |
| 6 | GCA_900102115.1 | *Blastococcus fimeti* DSM 44205$^T$ | DTEF |
| 7 | GCA_900230285.1 | *Blastococcus haudaquaticus* DSM 44270$^T$ | DTEF |
| 8 | GCA_003075095.1 | *Blastococcus litoris* GP-S2-8$^T$ | |
| 9 | GCA_900188025.1 | *Blastococcus mobilis* DSM 44272$^T$ | DTEF |
| 10 | GCA_004217455.1 | *Blastococcus saxobsidens* DSM 44509$^T$ | |
| 11 | GCA_900112815.1 | *Blastococcus tunisiensis* DSM 46838$^T$ | |
| 12 | GCA_008124835.1 | *Blastococcus xanthinilyticus* DSM 46842$^T$ | |
| 13 | GCA_900107105.1 | *Geodermatophilus africanus* DSM 45422$^T$ | DTEF |
| 14 | GCA_900116515.1 | *Geodermatophilus amargosae* DSM 46136$^T$ | DTEF |
| 15 | GCA_900182545.1 | *Geodermatophilus aquaeductus* DSM 46834$^T$ | |
| 16 | GCA_016907675.1 | *Geodermatophilus bullaregiensis* DSM 46841$^T$ | DTEF |
| 17 | GCA_002802985.1 | *Geodermatophilus chilensis* B12$^T$ | |
| 18 | GCA_013408985.1 | *Geodermatophilus daqingensis* DSM 104001$^T$ | |
| 19 | GCA_900115395.1 | *Geodermatophilus dictyosporus* DSM 43161$^T$ | DTEW, DTEF |
| 20 | GCA_003428645.1 | *Geodermatophilus marinus* LHW52908$^T$ | HeR |
| 21 | GCA_900129495.1 | *Geodermatophilus nigrescens* DSM 45408$^T$ | DTEF |
| 22 | GCA_003182485.1 | *Geodermatophilus normandii* DSM 45417$^T$ | DTEF |
| 23 | GCA_000025345.1 | *Geodermatophilus obscurus* DSM 43160$^T$ | DTEF |
| 24 | GCA_900111455.1 | *Geodermatophilus poikilotrophus* DSM 44209$^T$ | DTEF |
| 25 | GCA_900188375.1 | *Geodermatophilus pulveris* DSM 46839$^T$ | DTEW, DTEF, HeR |
| 26 | GCA_900114385.1 | *Geodermatophilus ruber* DSM 45317$^T$ | |
| 27 | GCA_014191795.1 | *Geodermatophilus sabuli* CECT 8820$^T$ | DTEF |
| 28 | GCA_900188205.1 | *Geodermatophilus saharensis* DSM 45423$^T$ | DTEF |
| 29 | GCA_900103785.1 | *Geodermatophilus siccatus* DSM 45419$^T$ | DTEF |
| 30 | GCA_900102745.1 | *Geodermatophilus telluris* DSM 45421$^T$ | DTEF |
| 31 | GCA_003002915.1 | *Geodermatophilus tzadiensis* DSM 45416$^T$ | DTEF |

| | | | |
|---|---|---|---|
| 32 | GCA_010685995.1 | *Goekera deserti* CPCC 205119$^T$ | DTEW |
| 33 | GCA_900100695.1 | *Klenkia brasiliensis* DSM 44526$^T$ | |
| 34 | GCA_900101025.1 | *Klenkia marina* DSM 45722$^T$ | DTEF |
| 35 | GCA_900103975.1 | *Klenkia soli* DSM 45843$^T$ | |
| 36 | GCA_900112525.1 | *Klenkia taihuensis* DSM 45962$^T$ | DTEF |
| 37 | GCA_005930475.1 | *Modestobacter altitudinis* 1G4$^T$ | DTEF |
| 38 | GCA_000761485.1 | *Modestobacter caceresii* KNN 45-2b$^T$ | NDQ, HeR[2] |
| 39 | GCA_005930495.1 | *Modestobacter excelsi* 1G6$^T$ | DTEF |
| 40 | GCA_000306785.1 | *Modestobacter italicus* BC501$^T$ | |
| 41 | GCA_019779905.1 | *Modestobacter lapidis* DSM 100206$^T$ | |
| 42 | GCA_011758655.1 | *Modestobacter marinus* DSM 45201$^T$ | NDQ, HeR |
| 43 | GCA_010686625.1 | *Modestobacter muralis* DSM 100205$^T$ | DTEF |
| 44 | GCA_007994135.1 | *Modestobacter roseus* DSM 45764$^T$ | NDQ, HeR |
| 45 | GCA_014195485.1 | *Modestobacter versicolor* DSM 16678$^T$ | |
| 46 | GCA_900110555.1 | *Trujillonella endophytica* DSM 45413$^T$ | |
| 47 | GCA_010682065.1 | *Blastococcus* sp. 67B17 | DTEF |
| 48 | GCA_000582785.1 | *Blastococcus* sp. AP3 | |
| 49 | GCA_004570425.1 | *Blastococcus* sp. AZ43 | DTEF |
| 50 | GCA_030731085.1 | *Blastococcus* sp. BMG 814 | |
| 51 | GCA_005222955.1 | *Blastococcus* sp. CCUG 61487 | |
| 52 | GCA_019800035.1 | *Blastococcus* sp. CPCC 205347 | |
| 53 | GCA_004570875.1 | *Blastococcus* sp. CT_GayMR16 | |
| 54 | GCA_004570385.1 | *Blastococcus* sp. CT_GayMR19 | |
| 55 | GCA_004570345.1 | *Blastococcus* sp. CT_GayMR20 | |
| 56 | GCA_000284015.1 | *Blastococcus* sp. DD2 | |
| 57 | GCA_900109665.1 | *Blastococcus* sp. DSM 46786 | |
| 58 | GCA_031845325.1 | *Blastococcus* sp. DSM 46792 | HeR[2] |
| 59 | GCA_021698975.1 | *Blastococcus* sp. KM273128 | |
| 60 | GCA_021698995.1 | *Blastococcus* sp. KM273129 | |
| 61 | GCA_020166105.1 | *Blastococcus* sp. LR1 | DTEF |
| 62 | GCA_021699055.1 | *Blastococcus* sp. MG754426 | |
| 63 | GCA_021699025.1 | *Blastococcus* sp. MG754427 | |
| 64 | GCA_023016265.1 | *Blastococcus* sp. PRF04-17 | |
| 65 | GCA_003319095.1 | *Blastococcus* sp. TBT05-19 | |

| | | | |
|---|---|---|---|
| 66 | GCA_003319175.1 | *Blastococcus* sp. TF02-8 | |
| 67 | GCA_003319105.1 | *Blastococcus* sp. TF02-9 | |
| 68 | GCA_004571075.1 | *Blastococcus* sp. TF02A_35 | DTEF |
| 69 | GCA_003319185.1 | *Blastococcus* sp. TF02A-30 | HeR |
| 70 | GCA_016612845.1 | *Blastococcus* sp. TML/C7B | |
| 71 | GCA_016612865.1 | *Blastococcus* sp. TML/M2B | |
| 72 | GCA_000620185.1 | *Blastococcus* sp. URHB0062 | |
| 73 | GCA_026967805.1 | *Blastococcus* sp. VKM Ac-2987 | NDQ |
| 74 | GCA_004571065.1 | *Geodermatophilus* sp. DF01_2 | |
| 75 | GCA_900143215.1 | *Geodermatophilus* sp. DSM 43162 | DTEF |
| 76 | GCA_900115505.1 | *Geodermatophilus* sp. DSM 44208 | DTEF |
| 77 | GCA_032460525.1 | *Geodermatophilus* sp. DSM 44513 | DTEW, NTQ, HeR |
| 78 | GCA_900104025.1 | *Geodermatophilus* sp. DSM 45219 | DTEF |
| 79 | GCA_010686745.1 | *Geodermatophilus* sp. I12A-02606 | DTEF |
| 80 | GCA_010686765.1 | *Geodermatophilus* sp. I12A-02694 | DTEF |
| 81 | GCA_003319115.1 | *Geodermatophilus* sp. TF02-6 | DTEF |
| 82 | GCA_025716945.1 | *Geodermatophilus* sp. YIM_151500 | |
| 83 | GCA_009296285.1 | *Goekera* sp. CPCC 205215 | DTEW |
| 84 | GCA_010077965.1 | *Goekera* sp. CPCC 205251 | DTEW |
| 85 | GCA_900491935.1 | *Klenkia* sp. IHUMI-26.3 | DTEW |
| 86 | GCA_001424455.1 | *Klenkia* sp. Leaf369 | DTEW |
| 87 | GCA_001424485.1 | *Klenkia* sp. Leaf380 | DTEF |
| 88 | GCA_014218215.1 | *Klenkia* sp. NBWT11 | DTEW |
| 89 | GCA_024124375.1 | *Klenkia* sp. PcliD-1-E | DTEF |
| 90 | GCA_019799965.1 | *Modestobacter* sp. CPCC 205242 | |
| 91 | GCA_019799945.1 | *Modestobacter* sp. CPCC 205245 | |
| 92 | GCA_900107175.1 | *Modestobacter* sp. DSM 44400 | |
| 93 | GCA_011039295.1 | *Modestobacter* sp. KNN46-3 | NDQ[3] |
| 94 | GCA_019112525.1 | *Modestobacter* sp. L9-4 | DTEF |
| 95 | GCA_000620205.1 | *Modestobacter* sp. URHA0031 | DTEW |
| 96 | GCA_001691715.2 | *Modestobacter* sp. VKM Ac-2676 | NDQ |
| 97 | GCA_026968045.1 | *Modestobacter* sp. VKM Ac-2977 | NDQ, HeR |
| 98 | GCA_026968025.1 | *Modestobacter* sp. VKM Ac-2978 | NDQ, HeR[2] |
| 99 | GCA_026967995.1 | *Modestobacter* sp. VKM Ac-2979 | NDQ, HeR |

| | | | |
|---|---|---|---|
| 100 | GCA_026967965.1 | *Modestobacter* sp. VKM Ac-2980 | NDQ, HeR |
| 101 | GCA_026967905.1 | *Modestobacter* sp. VKM Ac-2981 | NDQ, HeR |
| 102 | GCA_026967935.1 | *Modestobacter* sp. VKM Ac-2982 | NDQ, HeR |
| 103 | GCA_026967885.1 | *Modestobacter* sp. VKM Ac-2983 | DTEW, NDQ, HeR |
| 104 | GCA_026967985.1 | *Modestobacter* sp. VKM Ac-2984 | DTEF, NDQ, HeR |
| 105 | GCA_026967925.1 | *Modestobacter* sp. VKM Ac-2985 | DTEW, NDQ, HeR |
| 106 | GCA_026967795.1 | *Modestobacter* sp. VKM Ac-2986 | DTEW, DTEF |
| 107 | GCA_019799905.1 | *Trujillonella* sp. CPCC 205247 | DTEF |
| 108 | GCA_019799925.1 | *Trujillonella* sp. CPCC 205250 | DTEF |
| 109 | GCA_003319125.1 | *Trujillonella* sp. TF02A-26 | DTEF |
| 110 | GCA_000701365.1 | *Trujillonella* sp. URHD0036 | |

[1] DTEW, rhodopsins with aspartate, threonine, glutamate and tryptophan in positions 85, 89, 96 and 182 (in BR numbering), respectively; DTEF, rhodopsins with aspartate, threonine, glutamate and phenylalanine in positions 85, 89, 96 and 182 (in BR numbering), respectively; NDQ, rhodopsins with asparagine, aspartate and glutamine in positions 85, 89 and 96 (in BR numbering), respectively; NTQ, rhodopsins with asparagine, threonine and glutamine in positions 85, 89 and 96 (in BR numbering), respectively; HeR, heliorhodopsin.

[2] fragmented gene

[3] two identical genes

**Supplementary Table S2.** Rhodopsin genes found in members of the family *Geodermatophilaceae*.

| # | Assembly Acc. No. | Organism | Rhodopsin | Protein Acc. No. or Gene Location |
|---|---|---|---|---|
| 1 | GCA_900221005.1 | *Blastococcus aggregatus* DSM 4725$^{T}$ | HeR | OBQI01000003.1:233976-234767 |
| 2 | GCA_002938425.1 | *Blastococcus atacamensis* P6$^{T}$ | DTEF | POQU01000005.1:rc(97833-98786) |
| 3 | GCA_900102425.1 | *Blastococcus aurantiacus* DSM 44268$^{T}$ | DTEF | SDF05506.1 |
| 4 | GCA_030165005.1 | *Blastococcus capsensis* BMG 804$^{T}$ | NDQ | MDK3256622.1 |
| 5 | GCA_900102115.1 | *Blastococcus fimeti* DSM 44205$^{T}$ | DTEF | SDE68305.1 |
| 6 | GCA_900230285.1 | *Blastococcus haudaquaticus* DSM 44270$^{T}$ | DTEF | SOD99373.1 |
| 7 | GCA_900188025.1 | *Blastococcus mobilis* DSM 44272$^{T}$ | DTEF | SNR30776.1 |
| 8 | GCA_900107105.1 | *Geodermatophilus africanus* DSM 45422$^{T}$ | DTEF | SDX60326.1 |
| 9 | GCA_900116515.1 | *Geodermatophilus amargosae* DSM 46136$^{T}$ | DTEF | SFT51832.1 |
| 10 | GCA_016907675.1 | *Geodermatophilus bullaregiensis* DSM 46841$^{T}$ | DTEF | MBM7808800.1 |
| 11 | GCA_900115395.1 | *Geodermatophilus dictyosporus* DSM 43161$^{T}$ | DTEW | SFO59867.1 |
| 12 | GCA_900115395.1 | *Geodermatophilus dictyosporus* DSM 43161$^{T}$ | DTEF | SFN82483.1 |
| 13 | GCA_003428645.1 | *Geodermatophilus marinus* LHW52908$^{T}$ | HeR | RFU19957.1 |
| 14 | GCA_900129495.1 | *Geodermatophilus nigrescens* DSM 45408$^{T}$ | DTEF | SHG30246.1 |
| 15 | GCA_003182485.1 | *Geodermatophilus normandii* DSM 45417$^{T}$ | DTEF | PWW23913.1 |
| 16 | GCA_000025345.1 | *Geodermatophilus obscurus* DSM 43160$^{T}$ | DTEF | CP001867.1:rc(3831631-3832608) |
| 17 | GCA_900111455.1 | *Geodermatophilus poikilotrophus* DSM 44209$^{T}$ | DTEF | SET23820.1 |
| 18 | GCA_900188375.1 | *Geodermatophilus pulveris* DSM 46839$^{T}$ | DTEW | SNS94839.1 |
| 19 | GCA_900188375.1 | *Geodermatophilus pulveris* DSM 46839$^{T}$ | DTEF | SNS80697.1 |
| 20 | GCA_900188375.1 | *Geodermatophilus pulveris* DSM 46839$^{T}$ | HeR | SNS83322.1 |

| | | | | |
|---|---|---|---|---|
| 21 | GCA_014191795.1 | *Geodermatophilus sabuli* CECT 8820$^{T}$ | DTEF | MBB3085333.1 |
| 22 | GCA_900188205.1 | *Geodermatophilus saharensis* DSM 45423$^{T}$ | DTEF | SNS20283.1 |
| 23 | GCA_900103785.1 | *Geodermatophilus siccatus* DSM 45419$^{T}$ | DTEF | SDM18943.1 |
| 24 | GCA_900102745.1 | *Geodermatophilus telluris* DSM 45421$^{T}$ | DTEF | SDD54308.1 |
| 25 | GCA_003002915.1 | *Geodermatophilus tzadiensis* DSM 45416$^{T}$ | DTEF | PRY49980.1 |
| 26 | GCA_010685995.1 | *Goekera deserti* CPCC 205119$^{T}$ | DTEW | NEL55319.1 |
| 27 | GCA_900101025.1 | *Klenkia marina* DSM 45722$^{T}$ | DTEF | SCX56670.1 |
| 28 | GCA_900112525.1 | *Klenkia taihuensis* DSM 45962$^{T}$ | DTEF | SFD31338.1 |
| 29 | GCA_005930475.1 | *Modestobacter altitudinis* 1G4$^{T}$ | DTEF | SJEW01000037.1:137-1027 |
| 30 | GCA_000761485.1 | *Modestobacter caceresii* KNN 45-2b$^{T}$ | NDQ | JPMX01000052.1:rc(34964-35803) |
| 31 | GCA_000761485.1 | *Modestobacter caceresii* KNN 45-2b$^{T}$ | HeR[1] | JPMX01000052.1:21695-22485 |
| 32 | GCA_005930495.1 | *Modestobacter excelsi* 1G6$^{T}$ | DTEF | SJEX01000014.1:rc(9608-10498) |
| 33 | GCA_011758655.1 | *Modestobacter marinus* DSM 45201$^{T}$ | NDQ | NIH68942.1 |
| 34 | GCA_011758655.1 | *Modestobacter marinus* DSM 45201$^{T}$ | HeR | NIH68948.1 |
| 35 | GCA_010686625.1 | *Modestobacter muralis* DSM 100205$^{T}$ | DTEF | JAAGWB010000013.1:rc(518948-519886) |
| 36 | GCA_007994135.1 | *Modestobacter roseus* DSM 45764$^{T}$ | NDQ | TWH72922.1 |
| 37 | GCA_007994135.1 | *Modestobacter roseus* DSM 45764$^{T}$ | HeR | TWH72921.1 |
| 38 | GCA_010682065.1 | *Blastococcus* sp. 67B17 | DTEF | NEK86822.1 |
| 39 | GCA_004570425.1 | *Blastococcus* sp. AZ43 | DTEF | TFV62144.1 |
| 40 | GCA_031845325.1 | *Blastococcus* sp. DSM 46792 | HeR[1] | JAVREI010000002.1:rc(256665-257033) |
| 41 | GCA_020166105.1 | *Blastococcus* sp. LR1 | DTEF | MCA0146817.1 |
| 42 | GCA_004571075.1 | *Blastococcus* sp. TF02A_35 | DTEF | SPQP01000030.1:rc(2242-3162) |

| | | | | |
|---|---|---|---|---|
| 43 | GCA_003319185.1 | *Blastococcus* sp. TF02A-30 | HeR | RBY84940.1 |
| 44 | GCA_026967805.1 | *Blastococcus* sp. VKM Ac-2987 | NDQ | MCZ2860642.1 |
| 45 | GCA_900143215.1 | *Geodermatophilus* sp. DSM 43162 | DTEF | SHN70418.1 |
| 46 | GCA_900115505.1 | *Geodermatophilus* sp. DSM 44208 | DTEF | SFO70549.1 |
| 47 | GCA_032460525.1 | *Geodermatophilus* sp. DSM 44513 | DTEW | WNV77007.1 |
| 48 | GCA_032460525.1 | *Geodermatophilus* sp. DSM 44513 | NTQ | WNV74680.1 |
| 49 | GCA_032460525.1 | *Geodermatophilus* sp. DSM 44513 | HeR | WNV77816.1 |
| 50 | GCA_900104025.1 | *Geodermatophilus* sp. DSM 45219 | DTEF | SDO47926.1 |
| 51 | GCA_010686745.1 | *Geodermatophilus* sp. I12A-02606 | DTEF | JAAGWE010000002.1:139297-140265 |
| 52 | GCA_010686765.1 | *Geodermatophilus* sp. I12A-02694 | DTEF | NEK56946.1 |
| 53 | GCA_003319115.1 | *Geodermatophilus* sp. TF02-6 | DTEF | QOHF01000008.1:100188-101198 |
| 54 | GCA_009296285.1 | *Goekera* sp. CPCC 205215 | DTEW | MPR00383.1 |
| 55 | GCA_010077965.1 | *Goekera* sp. CPCC 205251 | DTEW | NDI50414.1 |
| 56 | GCA_900491935.1 | *Klenkia* sp. IHUMI-26.3 | DTEW | SSC23996.1 |
| 57 | GCA_001424455.1 | *Klenkia* sp. Leaf369 | DTEW | LMQA01000001.1:rc(1544766-1545665) |
| 58 | GCA_001424485.1 | *Klenkia* sp. Leaf380 | DTEF | LMQC01000006.1:rc(39678-40544) |
| 59 | GCA_014218215.1 | *Klenkia* sp. NBWT11 | DTEW | QNG37172.1 |
| 60 | GCA_024124375.1 | *Klenkia* sp. PcliD-1-E | DTEF | MCO7222595.1 |
| 61 | GCA_011039295.1 | *Modestobacter* sp. KNN46-3 | NDQ[2] | JAAKGN010000001.1:63179-64018 |
| 62 | GCA_011039295.1 | *Modestobacter* sp. KNN46-3 | NDQ[2] | JAAKGN010000001.1:rc(135481-136320) |
| 63 | GCA_019112525.1 | *Modestobacter* sp. L9-4 | DTEF | CP077800.1:rc(2283185-2284093) |
| 64 | GCA_000620205.1 | *Modestobacter* sp. URHA0031 | DTEW | JHVP01000005.1:rc(245422-246336) |

| | | | | |
|---|---|---|---|---|
| 65 | GCA_001691715.2 | *Modestobacter* sp. VKM Ac-2676 | NDQ | OMQ16403.1 |
| 66 | GCA_026968045.1 | *Modestobacter* sp. VKM Ac-2977 | NDQ | MCZ2820408.1 |
| 67 | GCA_026968045.1 | *Modestobacter* sp. VKM Ac-2977 | HeR | MCZ2820399.1 |
| 68 | GCA_026968025.1 | *Modestobacter* sp. VKM Ac-2978 | NDQ | MCZ2848615.1 |
| 69 | GCA_026968025.1 | *Modestobacter* sp. VKM Ac-2978 | HeR[1] | JAPVBP010000004.1:118574-119366 |
| 70 | GCA_026967995.1 | *Modestobacter* sp. VKM Ac-2979 | NDQ | MCZ2811703.1 |
| 71 | GCA_026967995.1 | *Modestobacter* sp. VKM Ac-2979 | HeR | MCZ2811712.1 |
| 72 | GCA_026967965.1 | *Modestobacter* sp. VKM Ac-2980 | NDQ | MCZ2843426.1 |
| 73 | GCA_026967965.1 | *Modestobacter* sp. VKM Ac-2980 | HeR | MCZ2843435.1 |
| 74 | GCA_026967905.1 | *Modestobacter* sp. VKM Ac-2981 | NDQ | MCZ2826200.1 |
| 75 | GCA_026967905.1 | *Modestobacter* sp. VKM Ac-2981 | HeR | MCZ2826190.1 |
| 76 | GCA_026967935.1 | *Modestobacter* sp. VKM Ac-2982 | NDQ | MCZ2852735.1 |
| 77 | GCA_026967935.1 | *Modestobacter* sp. VKM Ac-2982 | HeR | MCZ2852745.1 |
| 78 | GCA_026967885.1 | *Modestobacter* sp. VKM Ac-2983 | DTEW | MCZ2805545.1 |
| 79 | GCA_026967885.1 | *Modestobacter* sp. VKM Ac-2983 | NDQ | MCZ2805551.1 |
| 80 | GCA_026967885.1 | *Modestobacter* sp. VKM Ac-2983 | HeR | MCZ2805566.1 |
| 81 | GCA_026967985.1 | *Modestobacter* sp. VKM Ac-2984 | DTEF | MCZ2818135.1 |
| 82 | GCA_026967985.1 | *Modestobacter* sp. VKM Ac-2984 | NDQ | MCZ2816131.1 |
| 83 | GCA_026967985.1 | *Modestobacter* sp. VKM Ac-2984 | HeR | MCZ2816140.1 |
| 84 | GCA_026967925.1 | *Modestobacter* sp. VKM Ac-2985 | DTEW | MCZ2839206.1 |
| 85 | GCA_026967925.1 | *Modestobacter* sp. VKM Ac-2985 | NDQ | MCZ2839203.1 |
| 86 | GCA_026967925.1 | *Modestobacter* sp. VKM Ac-2985 | HeR | MCZ2839193.1 |

| | | | | |
|---|---|---|---|---|
| 87 | GCA_026967795.1 | *Modestobacter* sp. VKM Ac-2986 | DTEW | MCZ2829154.1 |
| 88 | GCA_026967795.1 | *Modestobacter* sp. VKM Ac-2986 | DTEF | MCZ2827338.1 |
| 89 | GCA_019799905.1 | *Trujillonella* sp. CPCC 205247 | DTEF | JABGCI010000008.1:83723-84607 |
| 90 | GCA_019799925.1 | *Trujillonella* sp. CPCC 205250 | DTEF | JABGCG010000009.1:rc(199171-200055) |
| 91 | GCA_003319125.1 | *Trujillonella* sp. TF02A-26 | DTEF | QOHG01000001.1:2249-3133 |

[1] fragmented gene

[2] two identical genes

For rhodopsin designations, see Supplementary Table S1.

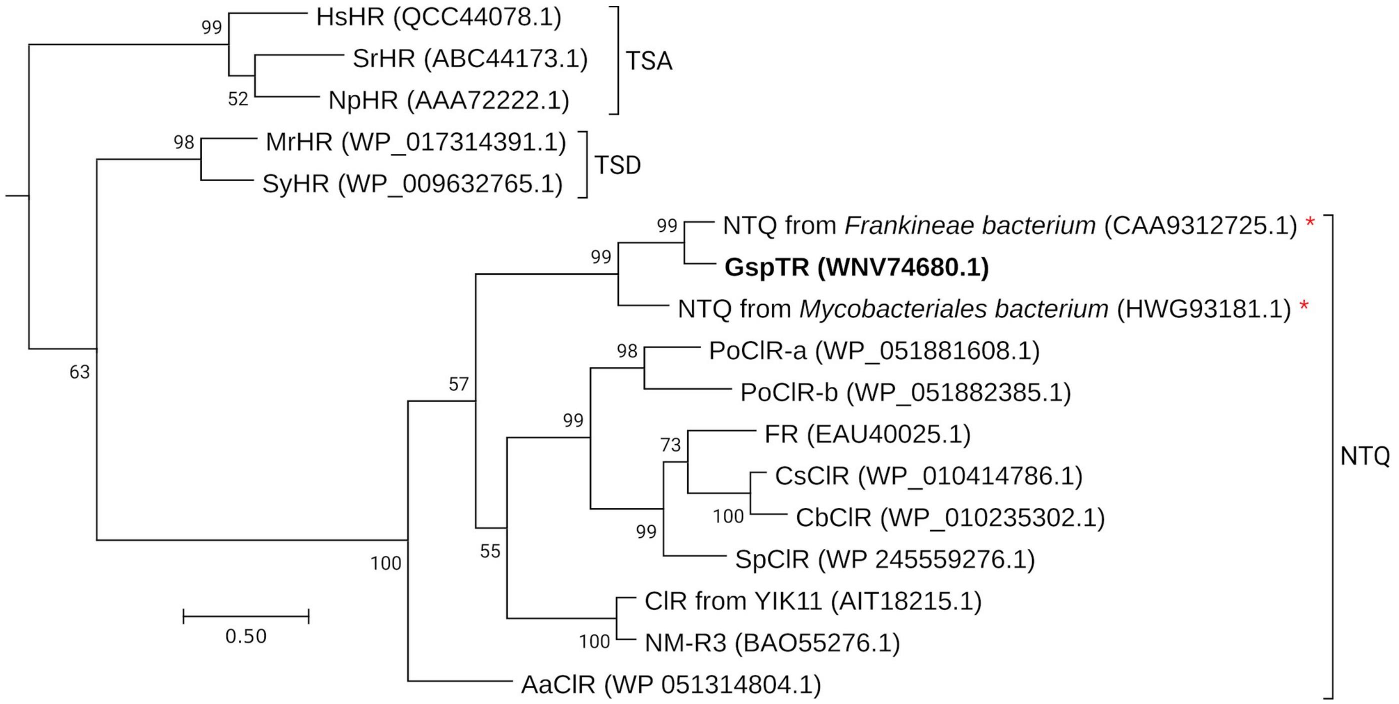


**Supplementary Fig. S3.** Maximum likelihood tree of protein sequences showing the relationship of GspTR rhodopsin from *Geodermatophilus* sp. DSM 44513 to the closest rhodopsins from actinomycetes and other previously characterized $Cl^-$-pumping rhodopsins. Sequence accession numbers for each protein are given in parentheses. Red asterisks denote the sequences closest to GspTR rhodopsin, available from the NCBI non-redundant protein database. The tree is drawn to scale, with branch lengths measured in the number of substitutions per site. Bootstrap values above 50% are indicated at the branch points. The sequence of bacteriorhodopsin (1QM8) served as an outgroup.

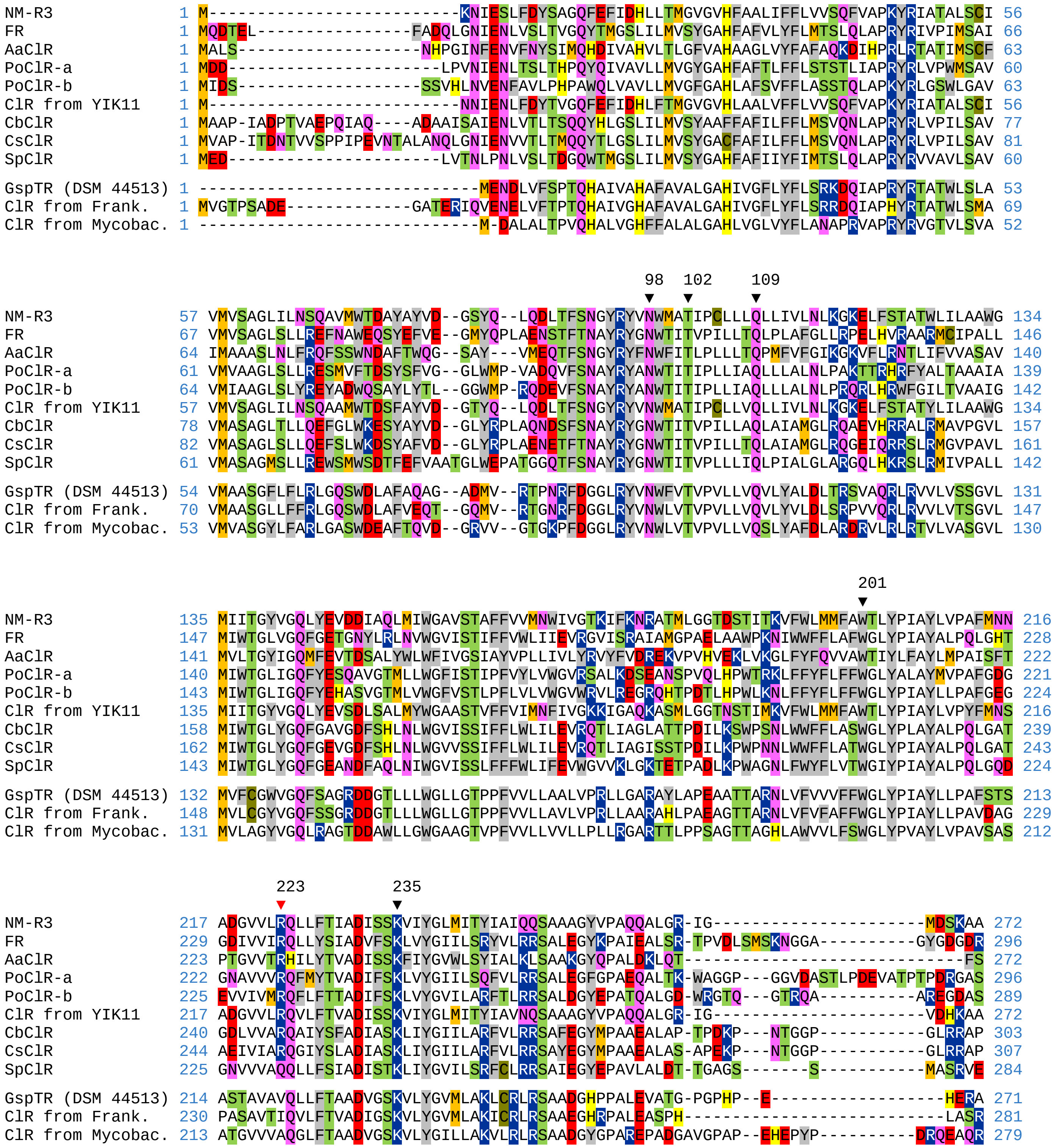


**Supplementary Figure S4.** Protein sequences alignment of the GspTR rhodopsin from *Geodermatophilus* sp. DSM 44513, the closest rhodopsins from actinomycetes and some previously characterized NTQ rhodopsins with Cl$^-$-pumping activity (ClR). Designations: NM-R3, ClR from *Nonlabens marinus* S1-08 (BAO55276.1); FR, ClR from *Fulvimarina pelagi* HTCC2506 (EAU40025.1); AaClR, ClR from *Alteribacter aurantiacus* DSM 18675 (WP_051314804.1); PoClR-a and PoClR-b, ClRs from *Parvularcula oceani* JLT2013 (WP_051881608.1 and WP_051882385.1, respectively); ClR from YIK11, ClR from *Nonlabens* sp. YIK11 (AIT18215.1); CbClR, ClR from *Citromicrobium bathyomarinum* JL354 (WP_010235302.1); CsClR, ClR from *Citromicrobium* sp. JLT1363 (WP_010414786.1); SpClR, ClR from *Novosphingopyxis baekryungensis* DSM 16222 (WP_245559276.1); ClR from Frank., NTQ rhodopsin from uncultured *Frankineae* (CAA9312725.1); ClR from Mycobac., NTQ rhodopsin from uncultured *Mycobacteriales* (HWG93181.1).

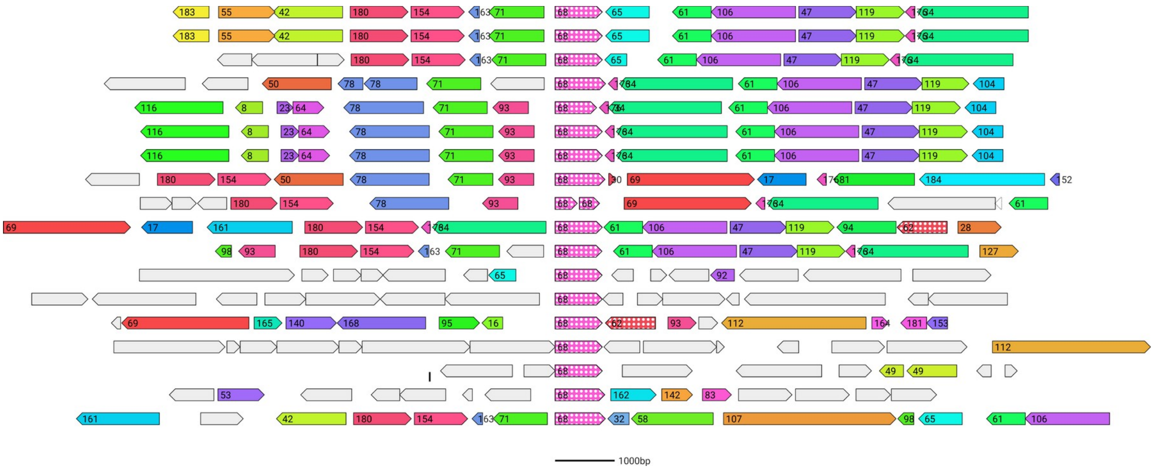


**Supplementary Figure S5.** Genomic neighbors of heliorhodopsin genes in *Geodermatophilaceae* strains.

```
BagHR (VKM Ac-1606) 1 GTGACCAGGCACACCCGTTCCGGGCCCGCACCGG-TCGCGACGGGGGTGGACGACGGGCGCCTTGCCGGCCTCCGGCGCT 79
HeR (KNN 45-2b)     1 GTCAGCGAGCTCGACAGTCCAGGATCCGCACCGG-TCGCGACCGGGGTGGACGACGCGCGCCTGGCCGGCCTCCGGCGCT 79
HeR (VKM Ac-2978)   1 GTGAGCGAGCTCGGCAGTCCAGGACCTGCACCGGGCCGCGACCGGGGTCGACGACGCGCGCCTGGCTGGCCTCCGGCGCT 80
HeR (DSM 46792)     0 -------------------------------------------------------------------------------- 0
HeR (g267)          1 ATGAGCGCGCTCGACAGTCCAGGACCCGCACCGG-TCGCGACCGGGGTGGACGACGCCCACCTGGCCGGCCTCCGGCGCT 79

BagHR (VKM Ac-1606) 80 GGAACCTGGCGCTGACGGTGCTGCATGTGGCCCAGGCCGTAGCGGTCCTGCTGCTGGCCAGCGGCTTCGCGATCCCCGTG 159
HeR (KNN 45-2b)     80 GGAACCTGGCGCTGACGGTGCTGCACGCAGCGCAGGCCATCGCCGCCCTGCTGCTGGCGAGCAGCTTCGCCCTCACCGTG 159
HeR (VKM Ac-2978)   81 GGAACCTGGCGCTGACCGTGCTGCACGCCGCCCAGGCCGCCGCCGTCCTGCTGCTGGCGAGCAGCTTCGCCATCACCGTG 160
HeR (DSM 46792)      0 -------------------------------------------------------------------------------- 0
HeR (g267)          80 GGAACCTGGCACTGACCGTGCTGCACGCCGCCCAGGCAGTCGCCGTCCTGCTGCTGGCGAGCAGCTTCGCCATCACCGTG 159

BagHR (VKM Ac-1606) 160 ACGTCGTCGTTCCCGGCCGGTCCGCCCGGGACTGCGGTACCGGCGCCGGAAGCCCTGGTCGACGTCGGCGTGGGGGCCGC 239
HeR (KNN 45-2b)     160 ACGTCGTCGTTCCCGGCCGGTCCCCCCGGGACGGCGGTGCCGGCACCGGAGCCGCTGGTCGACGTCGCGGTGGGCGTGGC 239
HeR (VKM Ac-2978)   161 ACGACGTCCTTCCCGGCCGGTCCGCCAGGCACGGCGGTGCCGGCGCCGGAGCCCCTGGTCGACATCGGGGTGGGGGCCGC 240
HeR (DSM 46792)       0 -------------------------------------------------------------------------------- 0
HeR (g267)          160 ACGACGTCCTTCCCGGCCGGTCCCCCCGGCACGGCGGTGCCGGCCCCGGAACCGCTGGTCGACATCGGCGTGGGGGTGGC 239

BagHR (VKM Ac-1606) 240 GATCGCGGTCTTCCTCGGGCTTGCCGCGCTGGACCACCTGCTGACCGCCACGGTGCTGCGCCGGCGTTACGAGGAGGACC 319
HeR (KNN 45-2b)     240 GATCGCGCTCTTCCTCGCGCTGGCCGCGGTGGACCACCTGCTGACCGCCACGGTGCTGCGCCGGCGCTACGAGGGCGACC 319
HeR (VKM Ac-2978)   241 CATCGCGGTCTTCCTCGGGCTCGCCGCGGTGGACCACCTGCTGACCGGCACGGTGCTCCGGCGGCGCTACGAGACCGACC 320
HeR (DSM 46792)       0 -------------------------------------------------------------------------------- 0
HeR (g267)          240 CATCGCGATCTTCCTCGGGCTGGCCGCGGTGGACCACCTGCTGACCGGCACCGTGCTCCGGCGCCGCTACGAGGACGACC 319

BagHR (VKM Ac-1606) 320 TGCGCCGCGGCATCAACCGGTTCCGCTGGCTGGAGTACTCGGTCAGCTCGACGATCATGGTGCTGCTGATCTGCGCCTAC 399
HeR (KNN 45-2b)     320 TGCGCCGTGGCATCAACCGGTTCCGCTGGGTCGAGTACTCGGTCAGCTCAACCATCATGGTGCTGCTGATCTGCGCCTAC 399
HeR (VKM Ac-2978)   321 TGCGCCGCGGCATCAACCGGTTCCGCTGGGTCGAGTACTCGGTCAGCTCCACCATCATGGTGCTGCTGATCTGCACCTAC 400
HeR (DSM 46792)       0 -------------------------------------------------------------------------------- 0
HeR (g267)          320 TGCGTCGCGGCATCAACCGGTTCCGCTGGTCGGAGTACTCGATCAGCTCGACGATCATGGTCCTGCTGATCTGCACCTAC 399

BagHR (VKM Ac-1606) 400 ACCGGCATCACCGGGCTGAGCGCGCTGATCGGCATCGCCGGTGCCAACGTCGCGATGATCCTCTTCGGTTGGCTGCAGGA 479
HeR (KNN 45-2b)     400 ACCGGCA-CACCGGGCTGAGCGCACTCATCGGCATCGCCGGCGCCAACGTCGCGATGATCCTCTTCGGCTGGTTGCAAGA 478
HeR (VKM Ac-2978)   401 ACCGGCATCACCGGGCTGAGCGCACTCATCGGCATCGCCGGCGCCAACGTCGCGATGATCCTCTTCGGCTGGTTGCAGGA 480
HeR (DSM 46792)       1 ------------------------CTCATCGGCATCGCCGGCGCCGACGTCGCCATGATCCTGTTCGGCTGGCTGCAGGA 56
HeR (g267)          400 ACCGGCATCACCGGACTGAGCGCACTGATCGGCATCGCCGGGGCGAACCTCGCGATGATCCTCTTCGGCTGGCTGCAGGA 479

BagHR (VKM Ac-1606) 480 GCTGATGAACCCGCCCGGCCGCACCCGGACGACCATGCAGCCGTTCTGGTTCGGCTGCGTTGCCGGGGCGGCGCCCTGGG 559
HeR (KNN 45-2b)     479 GCTGATGAACCCGCCCGGCCGCACCCGGACGACGATGCTGCCGTTCTGGTTCGGTTGCGTCGCCGGAGCCGCGCCCTGGG 558
HeR (VKM Ac-2978)   481 ACGGATGAACCCGCCCGGCCGCACCCGGACGACGATGCTGCCGTTCTGGTTCGGGTGCGTCGCCGGAGCCGCGCCCTGGG 560
HeR (DSM 46792)      57 AGTGATGAATCCGCCCGGCCGGGCCCGGACGACGATGCTGCCGTTCTGGTTCGGTTGCGTCGCCGGGGCTGCGCCGGGGG 136
HeR (g267)          480 GCGGATGAACCCGCCCGGCCGGACCCGGACGACGATGCTGCCGTTCTGGTTCGGGTGCATCGCCGGCGCCGCGCCCTGGA 559

BagHR (VKM Ac-1606) 560 TGGCGGTCACCGCCAACATCGTCGGGGCCGAGCAGATCCCCGGCTTCGTCTACGGGATCTTCGGATCGCTCTTCGTGTTC 639
HeR (KNN 45-2b)     559 TGGCGATCACCGCCAACGTCCTCGGGGCCGAGCAGCTACCCGGTTTCGTGCTCGGGATCTTCGGATCGCTGTTCGTCTTC 638
HeR (VKM Ac-2978)   561 TGGCGATCACCGCCAACGTCCTCGGGGCCGCGCAGATCCCCGGTTTCGTCCTCGGGATCTTCGGATCGCTGTTCGTCTTC 640
HeR (DSM 46792)     137 TGGCGGTCACGACCAACATCGTCGGCGCCGCGGAGATACCGGGCTTCGTCTACGGGATCTTCGGGTCGCTGTCCGTGTTC 216
HeR (g267)          560 TGGCCATCACCGCCAACATCGTCGGCGCCGCTCAGCTCCCGGGCTTCGTCGTCGGGATCTTCGTGTCGCTGTTCGTGTTC 639

BagHR (VKM Ac-1606) 640 TTCGCGAGCTTTGCGCTCAACCA----GTGGCTGCAGTACCGGGAGATCGGCCCGTGGCGCAGTTACGCCTACGGCGAGA 715
HeR (KNN 45-2b)     639 TTCATGAGCTTCGCGGTCAACCA----GTGGCTGCAGTACCGGGAGATCGGCCCGTGGCGCAGCTACGCCTACGGCGAGC 714
HeR (VKM Ac-2978)   641 TTCATGAGCTTCGCGGTCAACCA----GTGGCTGCAGTACCGGGAGATCGGCCCGTGGCGCAGCTACGCCTACGGCGAGC 716
HeR (DSM 46792)     217 TTCACGAGCTTCGCGGTCAACCA----GTGGCTGCAGTACCGGCAGGTCGGCGCGTGGCGCGACTACGCCTACGGCGAGC 292
HeR (g267)          640 TTCATGAGCTTCGCGGTCAACCAACCAGTGGCTGCAGTACCGGGAGATCGGCCCGTGGCGCAGCTACGCCTACGGCGAGA 719

BagHR (VKM Ac-1606) 716 AGGCCTACCTGGTGCTGAGCCTGGTCGCCAAGAGCGCCCTCGCCTGGCAGATCTTCGCCGGCTCGCTGGCCGGCTGA 792
HeR (KNN 45-2b)     715 GGGGGTACCTGGTGCTGAGCCTGGTCGCCAAGAGCCTGCTCGCCTGGCAGGTCTTCGCCGGCTCGCTGGCCACCTGA 791
HeR (VKM Ac-2978)   717 GGGGCTACCTGGTGCTGAGCCTGGTCGCCAAGAGCGCGCTCGCCTGGCAGGTCTTCGCCGGCTCGCTGGCCACCTGA 793
HeR (DSM 46792)     293 AGGCCTACCTGGTGCTCAGCCTGGTCGCCAAGAGCGCCCTCGCCTGGCAGGTCTTCGCCGGCTCGCTGGCCACCTGA 369
HeR (g267)          720 AGGCCTACCTCGTGCTGAGCCTGGTCGCCAAGAGCCTGCTCGCCTGGCAGGTCTTCGCCGGCTCACTGGCCACCTGA 796
```

**Supplementary Figure S6.** Nucleotide sequences alignment of heliorhodopsin pseudogenes. For designations, see Table 1 and Supplementary Table S1. Gene of BagHR used as reference sequence. g267, pseudogene from *Modestobacter* sp., unpublished genome. Insertion–deletion mutations are marked in red.

```
                                                                       23
                                                                       ▼
48C12               1 ------------------MAKP----------T--VKEIKSLQNFNRIAGVFHLLQMLAVLALANDFALPMTGTYLNGPP 50
HULAa30F3           1 ------------------MATD----------SATEKKLKGLRTWNIVVGLILAVQAVLIAVLTNSFALPVTATFMEGPP 52

BagHR (VKM Ac-1606) 1 MARR----GRHVPIQEVAVTRHTRSGPAPVATGVDDGRLAGLRRWNLALTVLHVAQAVAVLLLASGFAIPVTSSFPAGPP 76
HeR (DSM 46839)      1 ------------------MTGTTRPRSAPVATGVDDARLAGLRRWNLVLAVLHAAQAVAVLLLASDFAITVTSSFPAGPP 62
HeR (TF02A-30)      1 ------------------MTS-TGPGSAPVATGVDDARLAGLRRWNLALTVLHAAQAVAVLLLASSFAITVTSSFPAGPP 61
HeR (LHW52908)      1 ------------------MTRTAAPAPAPVATGVDDARLAGLRRWNLALAVLHAGQAVAVLVLASGFAITVTSSFPAGPP 62
HeR (DSM 45764)      1 ------------------MNRSTDGAPIPVATGVDDARLAGLRRWNLALTVLHAAQAIAVLLLASSFAITVTSSFPAGPP 62
HeR (DSM 44513)      1 ------------------MSGTALDRPAAVATGVDDARLAGLQRWNLALTVLHAAQAAAVLVLASGFAITVTSSFPAGPP 62
HeR (DSM 45201)      1 ------------------MSKPDSPGVVPVATGVDGARLAGLRRWNLALTVLHAAQTVAVVLLASSFAITVTTSFPTGPP 62
HeR (VKM Ac-2980)    1 ------------------MSELDSPGPVPAATGVDDARLAGLRRWNLALTVLHAAQAVAVLLLASSFAVTVTTSFPAGPP 62
HeR (VKM Ac-2983)    1 MLGPTADSGRP-STQEVVVSEPDSPGPAPVATGVDDARLAGLRRWNLALTVLHAAQAVAVLLLASSFAITVTTSFPAGPP 79
HeR (VKM Ac-2979)    1 ------------------MSELDSPGPVPAATGVDDARLAGLRRWNLALTVLHAAQAVAVLLLASSFAVTVTTSFPAGPP 62
HeR (VKM Ac-2977)    1 MVGCRKGSGPVLPDQEVVVTELDSPGSAPAATGVDDARLAGLRRWNLALTVLHAAQAIAVLLLASSFALTVTSSFPAGPP 80
HeR (VKM Ac-2985)    1 ------------------MSKPDSAGPAPVATGVDDALLAGLRRWNLALTVLHAAQAVAVLLLASSFAITVTTSFPTGPP 62
HeR (VKM Ac-2982)    1 ------------------MSEPDSPGSAPVATGVDDARLAGLRRWNLALTVLHAAQAVAVLLLASSFAITVTTSFPAGPP 62
HeR (VKM Ac-2984)    1 ------------------MSEPDSP--RPVATGVDDARLAGLRRWNLALTVLHAAQAVAVLLLASSFAITVTTSFPAGPP 60
HeR (VKM Ac-2981)    1 ------------------MSEPDSPGSAPVATGVDDARLAGLRRWNLALTVLHAAQAVAVLLLASSFAITVTTSFPAGPP 62

                                             76  80
                                             ▼   ▼
48C12               51 GTTFSAPVVILETPVGLAVALFLGLSALFHFIVSSGNFFKRYSASLMKNQNIFRWVEYSLSSSVMIVLIAQICGIADIVA 130
HULAa30F3           53 GTA-PALQHLFNIQTGWGVFIFMAISAGALLLIASPMVFSWYKRNLLESRNYGRWIEYFFSSSVMIVLISQICGISDIAA 131

BagHR (VKM Ac-1606) 77 GTAVPAPEALVDVGVGAAIAVFLGLAALDHLLTAT-VLRRRYEEDLRRGINRFRWLEYSVSSTIMVLLICAYTGITGLSA 155
HeR (DSM 46839)     63 GTPVPAPEPLVDVGVGAAIAVFLALAAVDHAVTAT-VGRRRYEEDLRGGLNRFRWLEYSLSSTVMVLLICAYTGITGLAA 141
HeR (TF02A-30)      62 GTAVPAPEPLVDVGVGAAIAVFLGLAALDHLLTAT-VLRRRYEEDLRRGLNRFRWLEYSVSSTIMVLLICAYTGITGLSA 140
HeR (LHW52908)      63 GTAVPAPEPLVDVGVGAAIAVFLGLAAVDHLVTAT-VLRRRYEEDLRRGINRFRWLEYSLSATIMVLLICAYTGITGLAA 141
HeR (DSM 45764)     63 GTAVPAPEPLVDVGVGAAIAVFLGLAALDHLLTGA-VLRRRYEEDLRGGINRFRWLEYSLSSTVMVLLICAYTGITGLSA 141
HeR (DSM 44513)     63 GAPVPAPEPLVDVGVGAAIAVFLALAAVDHLVTAT-VARRRYEDDLRRGINRFRWLEYSVSSTVMVLLIAAYTGITGLSA 141
HeR (DSM 45201)     63 GTPPPAPEPLVDIGVGSAIAVFLGLAAVDHLLTAT-ALRGRYEAGLRRGINRFRWSEYSISSTIMVLLICSYTGITGLSA 141
HeR (VKM Ac-2980)   63 GTAVPAPEPLVDIGVGAAIAVFLGLAAVDHLLTGT-VLRRRYETDLRRGINRFRWVEYSVSSTIMVLLICGYTGITGLSA 141
HeR (VKM Ac-2983)   80 GTAVPAPEPLVDIGVGAAIAVFLALAAVDHLLTGT-VLRRRYEDDLRGGINRFRWSEYSISSTIMVLLICTYTGITGLSA 158
HeR (VKM Ac-2979)   63 GTAVPAPEPLVDIGVGAAIAVFLGLAAVDHLLTGT-VLRRRYETDLRRGINRFRWVEYSVSSTIMVLLICGYTGITGLSA 141
HeR (VKM Ac-2977)   81 GTAVPAPAPLVDLGVGVAIALFLALAAVDHLLTAT-VLRRRYEDDLRRGINRFRWVEYSVSSTIMVLLICGYTGITGLSA 159
HeR (VKM Ac-2985)   63 GTAVPAPEPLVDIGVGVAIAVFLALAAVDHLLTGT-VLRRRYEDDLRGGINRFRWSEYSISSTIMVLLICTYTGITGLSA 141
HeR (VKM Ac-2982)   63 GTAVPAPEPLVDIGVGAAIAVFLALAAVDHLLTGT-VLRRRYEDDLRGGINRFRWSEYSISSTIMVLLICTYTGITGLSA 141
HeR (VKM Ac-2984)   61 GTAVPAPEPLVDIGVGAAIAVFLGLAAVDHLLTGT-VLRRRYETDLRRGINRFRWSEYSISSTIMVLLICTYTGITGLSA 139
HeR (VKM Ac-2981)   63 GTAVPAPEPLVDIGVGAAIAVFLALAAVDHLLTGT-VLRRRYEDDLRGGINRFRWSEYSISSTIMVLLICTYTGITGLSA 141

48C12               131 LLAIFGVNASMILFGWLQEKYTQPKD--GDLLPFWFGCIAGIVPWIGLLIYVIAPGSTSDVAVPGFVYGIIISLFLFFNS 208
HULAa30F3           132 LLAIFGINASMILFGALQEKYEKPGK--PNLLSFWFGSFAGIIPWIAVLIYVVSPG--VAAAPPAFVYGIVVALFVFFNC 207

BagHR (VKM Ac-1606) 156 LIGIAGANVAMILFGWLQELMNPPGRTRTTMQPFWFGCVAGAAPWVAVTANIVG-----AEQIPGFVYGIFGSLFVFFAS 230
HeR (DSM 46839)     142 LIGIAGANVAMILFGWLQELVNPPGRTRTTMLPFWFGCVAGAAPWVAITANILG-----AEQIPGFVYGIFGSLFVFFTS 216
HeR (TF02A-30)      141 LIGIAGANVAMILFGWLQELMNPPGRTRTTMLPFWFGCVAGAAPWVAVTANIVG-----AEQIPGFVYAIFVSLFVFFMS 215
HeR (LHW52908)      142 LIGIAGANVAMILFGWLQERANPPGRAGTTMLPFWFGCIAGAAPWVAITANIVG-----AEQIPGFVYGIFVSLFAFFTS 216
HeR (DSM 45764)     142 LIGIAGANVAMILFGWLQEQANPPGRARTTMLPFWFGCVAGVAPWVAITATIVG-----AEQIPGFVYGIFVSLFVFFAS 216
HeR (DSM 44513)     142 LIGIAGANVAMILFGWVQERVNPPGRTRTTMLPFWFGCVAGAAPWVAITANILG-----AEQIPGFVYGIFGSLFVFFAS 216
HeR (DSM 45201)     142 LIGIAGANVAMILFGWLQELMNPPGRTRTTMLPFWFGCVAGAAPWVAITANIVG-----AAQLPGFVIGIFVSLFVFFMS 216
HeR (VKM Ac-2980)   142 LIGIAGANVAMILFGWLQERMNPPGRTRTTMLPFWFGCVAGAAPWVAITANVLG-----AAQIPGFVLGIFGSLFVFFMS 216
HeR (VKM Ac-2983)   159 LIGIAGANVAMILFGWLQERMNPPGRTRTTMLPFWFGCIAGAAPWVAITANIVG-----AAQLPGFVIGIFVSLFVFFMS 233
HeR (VKM Ac-2979)   142 LIGIAGANVAMILFGWLQERMNPPGRTRTTMLPFWFGCVAGAAPWVAITANVLG-----AAQIPGFVLGIFGSLFVFFMS 216
HeR (VKM Ac-2977)   160 LIGIAGANVAMILFGWLQELMNPPGRTRTTMLPFWFGCVAGAAPWVAITANVLG-----AAQIPGFVLGIFGSLFVLFMS 234
HeR (VKM Ac-2985)   142 LIGIAGANVAMILFGWLQERMNPPGRTRTTMLPFWFGCIAGAAPWVAITANIVG-----AAQLPGFVIGIFVSLFAFFMS 216
HeR (VKM Ac-2982)   142 LIGIAGANVAMILFGWLQERMNPPGRTRTTMLPFWFGCIAGAAPWVAITANIVG-----AAQLPGFVIGIFVSLFVFFMS 216
HeR (VKM Ac-2984)   140 LIGIAGANVAMILFGWLQERMNPPGRTRTTMLPFWFGCIAGAAPWVAITANIVG-----AAQLPGFVIGIFVSLFVFFMS 214
HeR (VKM Ac-2981)   142 LIGIAGANVAMILFGWLQERMNPPGRTRTTMLPFWFGCIAGAAPWVAITANIVG-----AAQLPGFVIGIFVSLFVFFMS 216
```

*(continued on next page)*

(continue)

```
                                                  241
                                                   ▼
48C12               209 FALVQYLQYKGKGKWSNYLRGERAYIVLSLVAKSALAWQIFSGTLIPA 256
HULAa30F3           208 FAVNMILQYKQIGPWKDYLFGEKVYIILSLTAKSLLAWMVFANVL-VL 254

BagHR (VKM Ac-1606) 231 FALNQWLQYREIGPWRSYAYGEKAYLVLSLVAKSALAWQIFAGSL-AG 277
HeR (DSM 46839)     217 FAVNQWLQYRQLGPWRSYAYGERAYLVLSLVAKSALAWQVFAGSL-AT 263
HeR (TF02A-30)      216 FAVNQWLQYREIGPWRSYAYGEKAYLLLSLVAKSLLAWQIFAGSL-AT 262
HeR (LHW52908)      217 FAVNQWLQYREVGPWRSYAYGEKAYLVLSLVAKSALAWQVFAGSL-AG 263
HeR (DSM 45764)     217 FAVNQWLQYREVGPWRSYAFGEQAYLVLSLVAKSALAWQVFAGSL-AT 263
HeR (DSM 44513)     217 FAVNQWLQYRAIGPWRSYAYGERAYLVLSLVAKSALAWQVFAGSL-AT 263
HeR (DSM 45201)     217 FAVNQWLQYREVGPWRSYAFGEKAYLVLSLVAKSVLAWQVFAGSL-AT 263
HeR (VKM Ac-2980)   217 FAVNQWLQYREIGPWRSYAYGERGYLVLSLVAKSALAWQVFAGSL-AT 263
HeR (VKM Ac-2983)   234 FAVNQWLQYREIGPWRSYAHGEKAYLVLSLVAKSVLAWQVFAGSL-AT 280
HeR (VKM Ac-2979)   217 FAVNQWLQYREIGPWRSYAYGERGYLVLSLVAKSALAWQVFAGSL-AT 263
HeR (VKM Ac-2977)   235 FAVNQWLQYREIGPWRSYAYGERGYLVLSLVAKSLLAWQVFAGSL-AT 281
HeR (VKM Ac-2985)   217 FAVNQWLQYREIGPWRSYAYGEKAYLVLSLVAKSVLAWQVFAGSL-AT 263
HeR (VKM Ac-2982)   217 FAVNQWLQYREIGPWRSYAHGEKAYLVLSLVAKSVLAWQVFAGSL-AT 263
HeR (VKM Ac-2984)   215 FAVNQWLQYREIGPWRSYAHGEKAYLVLSLVAKSVLAWQVFAGSL-AT 261
HeR (VKM Ac-2981)   217 FAVNQWLQYREIGPWRSYAHGEKAYLVLSLVAKSVLAWQVFAGSL-AT 263
```

**Supplementary Figure S7.** Protein sequences alignment of heliorhodopsins from *Geodermatophilaceae* strains and the previously characterized heliorhodopsins 48C12 from uncultured actinomycete (AVZ43932.1) and HULAa30F3 from uncultured *Dehalococcoidia* (QOV09072.1). For designations of *Geodermatophilaceae* strains and rhodopsins, see Table 1, Supplementary Table S1 and Supplementary Table S2.

| | 148 | 156 | 157 | 160 | 184 | 191 | 194 | 197 | 201 | 205 | 207 | 208 | 211 |
|---|---|---|---|---|---|---|---|---|---|---|---|---|---|
| | ▼ | ▼ | ▼ | ▼ | ▼ | ▼ | ▼ | ▼ | ▼ | ▼ | ▼ | ▼ | ▼ |
| XR | L | G | F | T | R | N | L | L | G | I | Y | M | M |
| PRO95 | N | G | A | S | A | N | I | L | T | G | Y | L | Y |
| GdiWR (VKM Ac-659) ×3 | E | G | T | T | E | M | L | V | G | I | Y | L | V |
| DTEW (URHA0031) | E | G | T | T | E | M | I | V | G | I | Y | L | V |
| DTEW (Leaf369) ×4 | E | G | T | T | E | M | I | V | G | I | Y | L | I |
| DTEW (CPCC 205215) ×3 | D | G | T | T | E | M | L | V | G | I | Y | L | I |
| DTEW (DSM 46839) | E | G | T | T | Q | M | L | V | G | I | Y | L | V |
| DTEW (VKM Ac-2985) | E | G | T | T | E | M | L | V | G | I | Y | L | I |
| GdiFR (VKM Ac-659) | D | G | L | T | A | N | I | L | G | L | Y | A | V |
| GobFR (Ac-658) ×25 | D | G | L | T | E | N | I | L | G | L | Y | A | V |
| DTEF (Leaf380) | S | G | L | T | E | N | I | L | G | L | Y | S | V |
| DTEF (1G4) ×2 | S | G | L | T | E | N | I | L | G | L | Y | A | V |
| DTEF (DSM 45722) | S | G | L | T | E | N | V | L | G | L | Y | A | V |
| DTEF (DSM 44268) | D | G | L | T | E | N | I | F | G | L | Y | A | V |
| DTEF (DSM 45219) ×2 | D | G | L | T | E | N | V | L | G | L | Y | A | V |
| DTEF (DSM 45962) | S | G | V | S | E | N | I | L | G | L | Y | A | V |
| DTEF (DSM 45408) | D | G | L | T | E | N | L | L | G | L | Y | A | V |
| DTEF (DSM 45423) | D | G | L | T | P | N | I | L | G | L | Y | A | V |
| DTEF (DSM 46839) | D | G | L | T | E | N | T | L | G | V | Y | A | V |
| DTEF (DSM 44270) | D | G | L | T | E | N | T | L | G | L | Y | A | V |
| DTEF (DSM 100205) ×2 | D | G | L | C | E | N | I | L | G | L | Y | A | V |
| DTEF (PcliD-1-E) | S | G | I | T | A | N | I | L | G | L | Y | A | V |
| DTEF (VKM Ac-2984) | D | G | C | T | E | N | I | L | G | L | Y | A | V |
| MspDR (VKM Ac-2978) ×15 | R | G | S | T | E | A | W | L | M | G | Y | L | A |
| NDQ (BMG 804) | R | G | S | T | E | G | W | L | M | G | Y | L | A |
| GspTR (DSM 44513) | D | G | L | T | E | N | F | V | G | I | Y | L | A |

**Supplementary Figure S8.** Amino acids of Type I rhodopsins involved in carotenoid antenna binding. Identical variants are omitted, number after × signs indicate number of identical variants. Designations: XR, xanthorhodopsin from *Salinibacter ruber* DSM 13855 (ABC44767.1); PRO95, $Na^{+}$-pumping rhodopsin from *Dokdonia* sp. PRO95 (AEX55013.1). For designations of *Geodermatophilaceae* strains and rhodopsins, see Table 1, Supplementary Table S1 and Supplementary Table S2.

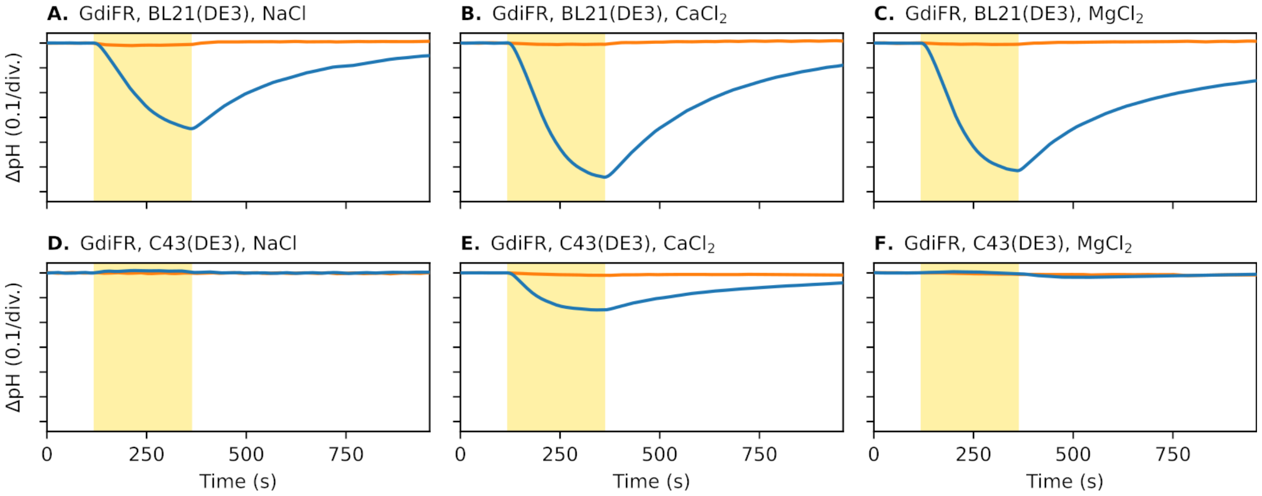


**Supplementary Figure S9.** Ion transport activity of DTEF rhodopsin from *G. dictyosporus* VKM Ac-659$^T$ (GdiFR). Measurements were performed with (orange line) and without (blue line) adding CCCP. The cell suspensions were illuminated for 240 s (yellow area on the plots).

```
                                                                                         61        74
KR2                     1 MTQELGNANFENFIGATEGFSEIAYQFTSHILTLGYAVMLAGLLYFILTIKNVDKKFQMSNILSAVVMVSAFLLLYAQAQ 80
NdR2                    1 MIQDLGNSNFENYVGATDGFSEMAYQMTSHVLTLGYAVMFAGLLYFILTIKNVDKKYRMSNILSAVVMVSAALLLYAQAG 80
PRO95                   1 MTQELGNANFENFIGATEGFSEIAYQFTSHILTLGYAVMLAGLLYFILTIKNVDKKFQMSNILSAVVMVSAFLLLYAQAQ 80
IAR                     1 MENL-GNATFENYIGLQDGFNEMAYQMVAHVLTLGYAVMLAGLFYFVLTIKTVAPRFRTSSVLSVVVMVSAFLLLYVQAS 79
GLR^x                   1 MTQELGNANFENFIGATEGFSEIAYQFTSHILTLGYAVMLAGLLYFILTIKKVDKKYQMSNILSAVVMVSAFLLLYAQAE 80
NM-R2                   1 MQQL-GNSNFENYIGASEGFSEMAYQMTSHVLTLGYAVMLAGLLYFILTIKKVDKRFQMSNILSAVVMVSAFLLLYAQAG 79

MspDR (VKM Ac-2978)     1 MN-------VEN----TFSYDFFQFTIVSHVFTFGYAAMAAGLVYFLLTSKNSLPRYQLSSTLSAVVMLSAFLELWVISQ 69
NDQ (DSM 101691)        1 MN-------VEN----TFSYDFFQFTIGSHVFTLGYAAMAAGLVYFLLTSKNSLPRYQLSSTLSAVVMLSAFLELLVISQ 69
NDQ (VKM Ac-2981)       1 MN-------VEN----TFTYDFFQYTIVSHVFTVGYAAMAAGLVYFVLTSKNSLPRYQLSSTLSAVVMVSAFLELLVISQ 69
NDQ (VKM Ac-2676)       1 MN-------VEN----TFTYDFFQYTIVSHVFTVGYAAMAAGLVYFVLTSKNSLPRYQLSSTLSAVVMVSAFLELLVISQ 69
NDQ (DSM 45764)         1 MN-------VEN----TFTYDFSQYTIVSHVFTLGYAAMAAGLVYFVLTSRNSLPRYRLSSTLSGVVMLSAFLELLVISQ 69
NDQ (DSM 45201)         1 MN-------VEN----TFTFDFFQYTIVSHVFTFGYAAMAAGLVYFVLTSKNSLPRYQLSSTLSAVVMLSAFLELLVISQ 69
NDQ (KNN46-3)           1 MN-------VEN----TFTYDFFQYTIVSHVFTLGYAAMAAGLVYFLLTSKNSLPRYQLSSTLSAVVMLSAFLELLVISL 69
NDQ (VKM Ac-2980)       1 MN-------VEN----TFSYDFFQFTIVSHVFTFGYAAMAAGLVYFLLTSKNSLPRYQLSSTLSAVVMLSAFLELWVISQ 69
NDQ (VKM Ac-2983)       1 MN-------VEN----TFTYDFFQYTIVSHVFTVGYAAMAAGLVYFVLTSKNSLPRYQLSSTLSAVVMVSAFLELLVISQ 69
NDQ (VKM Ac-2979)       1 MN-------VEN----TFSYDFFQFTIVSHVFTFGYAAMAAGLVYFLLTSKNSLPRYQLSSTLSAVVMLSAFLELWVISQ 69
NDQ (VKM Ac-2977)       1 MN-------VEN----TFSYDFFQFTIVSHVFTLGYAAMAAGLVYFLLTSKNSLPRYQLSSTLSAVVMLSAFLELLVTSQ 69
NDQ (VKM Ac-2987)       1 MN-------VEN----TFTYDFFQYTIVSHVFTLGYAAMAAGLVYFILTSKNSLPRYQLSSTLSAVVMLSAFLELLVISL 69
NDQ (VKM Ac-2982)       1 MN-------VEN----TFTYDFFQYTIVSHVFTVGYAAMAAGLVYFVLTSKNSLPRYQLSSTLSAVVMVSAFLELLVISQ 69
NDQ (VKM Ac-2984)       1 MN-------IEN----TFTYDFFQYTIVSHVFTVGYAAMAAGLVYFVLTSKNSLPRYQLSSTLSAVVMVSAFLELLVISQ 69
NDQ (VKM Ac-2985)       1 MN-------VEN----TFTYDFFQYTIVSHVFTVGYAAMAAGLVYFVLTSKNSLPRYQLSSTLSAVVMVSAFLELLVISQ 69
NDQ (BMG 804)           1 MN-------FEN----TFTYDFFQYTIVSHVFTLGYAAMAAGLAYFLLTSKNSLPRYQLSSTLSAVVMVSAFLELLVISL 69

                                                    112 116      123
KR2                    81 NWTSSFTFNEEVGRYFLDPSGDLFNNGYRYLNWLIDVPMLLFQILFVVSLTTSKFSSVRNQFWFSGAMMIITGYIGQFYE 160
NdR2                   81 NWTESFAFDAERGKYFLVEGGDLFNNGYRYLNWLIDVPMLLFQILFVVQLTKSKLSSVRNQFWFSGAMMIITGYIGQYYE 160
PRO95                  81 NWTSSFTYNEEVGRYFLDPSGDLFNNGYRYLNWLIDVPMLLFQILFVVSLTSSKFSSVRNQFWFSGAMMIITGYIGQFHE 160
IAR                    80 NWTESFVFDTERGKYFLGEGNDLFNNGYRYLNWLIDVPMLLFQILFVVTLTKSNFSSIRNQFWISGTGMIVTGYIGQFYE 159
GLR^x                  81 NWTTSFTFDISRGKYFLEPNGDLFNNGYRYLNWLIDVPMLLFQILFVVSLTKSKFSSIRNQFWFSGAMMIITGYIGQFYE 160
NM-R2                  80 NWTSSFTFDIELGRYFLDPDGDLFNNGYRYLNWLIDVPMLLFQILFVVTLTKSKLSSIRNQFWFSGTMMIITGYIGQFYE 159

MspDR (VKM Ac-2978)    70 RWDNAFSWDGEA---F-SSTGELFSNGYRYMNWTIDVPVLLLQMLIVLGVTGAAFRRGWIVFTVAGLGMIYTGYAGQFYE 145
NDQ (DSM 101691)       70 KWDSAFSWDGEA---F-SQTGTLFSNGYRYMNWTIDVPVLLLQMLIVLGVTGAAFRRGWIVFTVAGLGMIYTGYAGQFYE 145
NDQ (VKM Ac-2981)      70 KWESSFRWDGEA---F-SQADALFSNGYRYMNWSIDVPVLLLQMLIVLGVTGAAFRRGWIVFTVAGLGMVYTGYVGQFYE 145
NDQ (VKM Ac-2676)      70 KWESAFRWDGEA---F-SQTDALFSNGYRYMNWSIDVPVLLLQMLIVLGVTGAAFRRGWIVFTVAGLGMVYTGYVGQFYE 145
NDQ (DSM 45764)        70 RWDSAFRWDGEA---F-VQAGTLFSNGYRYMNWSIDVPVLLLQMLIVFGVTGAAFRRGWIVFTVAGLGMIYTGYVGQFHE 145
NDQ (DSM 45201)        70 KWDSAFSWDGEA---F-SPTGTLFSNGYRYMNWTIDVPVLLLQMLIVLGVTGAAFRRGWIVFTVAGLGMIYTGYAGQFYE 145
NDQ (KNN46-3)          70 RWDSAFSWDGEA---F-SSTGALFSNGYRYMNWTIDVPVLLLQMLIVLGVTGAAFRRGWIVFTVAGLGMIYTGYAGQFYE 145
NDQ (VKM Ac-2980)      70 RWDNAFSWDGEA---F-SSTGELFSNGYRYMNWTIDVPVLLLQMLIVLGVTGAAFRRGWIVFTVAGLGMIYTGYAGQFYE 145
NDQ (VKM Ac-2983)      70 KWESSFRWDGES---F-SQADALFSNGYRYMNWSIDVPVLLLQMLIVLGVTGAAFRRGWIVFTVAGLGMVYTGYVGQFYE 145
NDQ (VKM Ac-2979)      70 RWDNAFSWDGEA---F-SSTGELFSNGYRYMNWTIDVPVLLLQMLIVLGVTGAAFRRGWIVFTVAGLGMIYTGYAGQFYE 145
NDQ (VKM Ac-2977)      70 KWDSAFSWDGEA---F-SPTGTLFSNGYRYMNWTIDVPVLLLQMLIVLGVTGAAFRRGWIVFTVAGLGMIYTGYAGQFYE 145
NDQ (VKM Ac-2987)      70 RWDGAFTWDGEA---F-SSTGALFSNGYRYMNWTIDVPVLLTQLLIVLGITGAAFRRGWITFTVAGLAMIYTGYAGQFYE 145
NDQ (VKM Ac-2982)      70 KWESSFRWDGEA---F-SQADALFSNGYRYMNWSIDVPVLLLQMLIVLGVTGAAFRRGWIVFTVAGLGMVYTGYVGQFYE 145
NDQ (VKM Ac-2984)      70 KWESAFRWDGAA---F-SQADALFSNGYRYMNWSIDVPVLLLQMLIVLGVTGAAFRRGWIVFTVAGLGMVYTGYVGQFYE 145
NDQ (VKM Ac-2985)      70 KWESAFRWDGEA---F-SQTDALFSNGYRYMNWSIDVPVLLLQMLIVLGVTGAAFRRGWIMFTVAGLGMVYTGYVGQFYE 145
NDQ (BMG 804)          70 QWRSSFAWDGAA---Y-SQSAALFSNGYRYMNWSIDVPVLLTQLLIVLGVTGAAFRRGWITFTVAGLAMVYTGYVGQFYE 145

                                        180                                   215
KR2                   161 VSNLTAFLVWGAISSAFFFHILWVMKKVINEGKEGISPAGQKILSNIWILFLISWTLYPGAYLMPYLTGVDGFLYSEDGV 240
NdR2                  161 VTDLSAFFIWGAISTVFFFHILWLMNKVIKEGKVGIPKKGQKILSNIWILFLVSWFLYPGAYLMPHLGGIEGFLFNESGV 240
PRO95                 161 VSNLTAFLVWGAISSAFFFHILWVMKKVIKEGKEGISPAGQKILSNIWILFLISWTLYPGAYLMPYLTGVDGFLYSEDGV 240
IAR                   160 VTDLTMFAIWGAISTVFFFHILWLMKKVIDEGKDGIPAKAQETLQSIWVLFLVSWMLYPGAYLMPHLAGIEGLFFSEIGV 239
GLR^x                 161 VSNLTAFFVWGAISSVFFFHILWVMKKVINEGKEGLSADAQKILSNIWVLFLVSWFLYPGAYLMPYLTGLDGFFFSEDGV 240
NM-R2                 160 VSDITWFLIWGAISTVFFFHILYLMKKVINEGKEGISTKGQKILSNIWILFLISWFLYPGAYLMPYLGGIDGFLYNESGV 239

MspDR (VKM Ac-2978)   146 VERSAPFWIWGSISTVFFLVILALVYRTVYGNVGRLPAEVRPLVKAVWWLLLVSWMLYPGAYLMPAI------WDSADGV 219
NDQ (DSM 101691)      146 VERSAPFWIWGSISTVFFLVILALVYRTVYGNVGRLPAEVRPLVKAVWWLLLVSWMLYPGAYLMPAI------WDSADGV 219
NDQ (VKM Ac-2981)     146 VERSAPFWVWGSISTVFFLVILALVYRTVYGNVGRLPAEVRPLVKAVWWLLLVSWMLYPGAYLMPAI------WDSADGV 219
NDQ (VKM Ac-2676)     146 VERSAPFWIWGSISTVFFLVILALVYRTVYGNVGRLPAEVRPLVKAVWWLLLVSWMLYPGAYLMPAI------WDSADGV 219
NDQ (DSM 45764)       146 VERSAPFWVWGSISTVFFLVILALVYRTVYGNVGRMPAEVRPLVKAVWWLLLGSWMLYPGAYLMPAI------WDSADGV 219
NDQ (DSM 45201)       146 VERSAPFWIWGSISTVFFLVILVLVYRTVYGNVGRLPAEVRPLVKAVWWLLLVSWMLYPGAYLMPAI------WDSADGV 219
NDQ (KNN46-3)         146 VERSAPFWIWGSISTVFFLVILALVYRTVYGNVGRLPAEVRPLVKAVWWLLLVSWMLYPGAYLMPAI------WDSADGV 219
NDQ (VKM Ac-2980)     146 VERSAPFWIWGSISTVFFLVILALVYRTVYGNVGRLPAEVRPLVKAVWWLLLVSWMLYPGAYLMPAI------WDSADGV 219
NDQ (VKM Ac-2983)     146 VERSAPFWVWGSISTVFFLVILALVYRTVYGNVGRLPAEVRPLVKAVWWLLLVSWMLYPGAYLMPAI------WDSADGV 219
NDQ (VKM Ac-2979)     146 VERSAPFWIWGSISTVFFLVILALVYRTVYGNVGRLPAEVRPLVKAVWWLLLVSWMLYPGAYLMPAI------WDSADGV 219
NDQ (VKM Ac-2977)     146 VERSAPFWIWGSISTVFFLVILALVYRTVYGNVGRLPAEVRPLVKAVWWLLLVSWMLYPGAYLMPAI------WDSADGV 219
NDQ (VKM Ac-2987)     146 VERSAPFWVWGSISTVFFLVILWLVRRTVYGNVGRLPEEVRPLVKAVWWLLLVSWMLYPGAYLMPAL------WDSADGV 219
NDQ (VKM Ac-2982)     146 VERSAPFWVWGSISTVFFLVILALVYRTVYGNVGRLPAEVRPLVKAVWWLLLVSWMLYPGAYLMPAI------WDSADGV 219
NDQ (VKM Ac-2984)     146 VERSAPFWIWGSISTVFFLVILALVHRTVYGNVGRLPAEVRPLVKAVWWLLLVSWMLYPGAYLMPAI------WDSADGV 219
NDQ (VKM Ac-2985)     146 VERSAPFWVWGSISTVFFLVILALVYRTVYGNVGRLPAEVRPLVKAVWWLLLVSWMLYPGAYLMPAI------WDSADGV 219
NDQ (BMG 804)         146 VERSAPFWVWGSISTVFFLVLLVLVYRTIHGNLDRLPQEVRSLVRGVWWLLLFSWMLYPGAYLMPAM------WDSADGV 219
```

*(continued on next page)*

*(continue)*

```
                                           255     263
                                            ▼       ▼
KR2                 241 MARQLVYTIADVSSKVIYGVLLGNLAITLSKNKELVEA--------------------NS 280
NdR2                241 VGRQITYTIADVCSKVIYGVLLGNLALVLSKNKEMIE---------------------TA 279
PRO95               241 MARQLVYTIADVCSKVIYGVLLGNLAITLSKNKELVEA--------------------NS 280
IAR                 240 VARQITYTIADVSSKVIYGILLTNVAQVMSKEEGYLEH--------------------TT 279
GLRˣ                241 MARQLTYTIADVCSKVIYGVLLGNLALKLSNNKEMVEL--------------------SN 280
NM-R2               240 VGRQITYTIADVCSKVIYGVLLGNLAMTLSTKNKNEHK--------------------PA 279

MspDR (VKM Ac-2978) 220 VARQITYTVADVTSKVVYGILLAIIAQKVSKHEGHDAAVASDVTMPRGGITDVAAPGRAA 279
NDQ (DSM 101691)    220 VARQITYTVADITSKVVYGILLAVIARKVSKREGHDEAVASDVTMPRGGITDVAAPGRAA 279
NDQ (VKM Ac-2981)   220 VARQITYTVADVTSKVVYGILLAVIARKVSKHEGHDEAVASDVTMPRGGITEVAARGRAA 279
NDQ (VKM Ac-2676)   220 VARQITYTVADITSKVVYGILLAVIARKVSKHEGHDEAVASDVTLPRGGITEVAAPGRAA 279
NDQ (DSM 45764)     220 VARQITYTVADIVSKVVYGILLAVIAQKVSKHEHHEEALAADVTQPRGGITDVAAPGRAA 279
NDQ (DSM 45201)     220 VARQITYTIADVTSKVVYGILLAVIARKVSKHEGHDEAVASDVTMPRGGITDVAAPGRAA 279
NDQ (KNN46-3)       220 VARQITYTVADITSKVVYGILLAVIARKVSKHEGHDEAVASDVTMPRGGITDVAAPGRAA 279
NDQ (VKM Ac-2980)   220 VARQITYTVADVTSKVVYGILLAIIAQKVSKHEGHDAAVASDVTMPRGGITDVAAPGRAA 279
NDQ (VKM Ac-2983)   220 VARQITYTVADVTSKVVYGILLAVIARKVSKHEGHDEAVASDVTMPRGGITEVAAPGRAA 279
NDQ (VKM Ac-2979)   220 VARQITYTVADVTSKVVYGILLAIIAQKVSKHEGHDAAVASDVTMPRGGITDVAAPGRAA 279
NDQ (VKM Ac-2977)   220 VARQITYTVADVTSKVVYGILLAIIAQKVSKHEGHDAAVAADVTLPRGGITEVAAPGRAA 279
NDQ (VKM Ac-2987)   220 VARQITYTVADITSKVVYGILLAVIARKVSKHEGHDEAVAADVTMPRGGITEVAPPGRTT 279
NDQ (VKM Ac-2982)   220 VARQITYTVADVTSKVVYGILLAVIARKVSKHEGHDEAVASDVTMPRGGITEVAARGRAA 279
NDQ (VKM Ac-2984)   220 VARQITYTVADVTSKVVYGILLAVIARKVSKHEGHDEAIASDVTMPRGGITEVAAPGRAA 279
NDQ (VKM Ac-2985)   220 VARQITYTVADVTSKVVYGILLAVIARKVSKHEGHDEAVASDVTMPRGGITEVAAPGRAA 279
NDQ (BMG 804)       220 VARQITYTVADVTSKVVYGILLAAIARKISKHEGHDPATAADVTMPRGGVTDTAAPGR-I 278
```

**Supplementary Figure S10.** Protein sequences alignment of the NDQ rhodopsins from the *Geodermatophilaceae* family and some previously characterized NDQ rhodopsins with $H^+/Na^+$-, $Na^+$-pumping activity (NaR). Designations: KR2, NaR from *Dokdonia eikasta* NBRC 100814 (BAN14808.1); NdR2, NaR from *Nonlabens dokdonensis* DSW-6 (AGC76155.1); PRO95, NaR from *Dokdonia* sp. PRO95 (AEX55013.1); IAR, NaR from *Indibacter alkaliphilus* LW1 (EOZ93469.1); GLR, NaR from *Gillisia limnaea* DSM 15749 (EHQ02967.1); NM-R2, NaR from *Nonlabens marinus* S1-08 (BAO54106.1). For designations of *Geodermatophilaceae* strains and rhodopsins, see Table 1, Supplementary Table S1 and Supplementary Table S2. ˣ, 61 amino acids were skipped from N-terminus.
